# Increased spontaneous EEG signal diversity during stroboscopically-induced altered states of consciousness

**DOI:** 10.1101/511766

**Authors:** David J. Schwartzman, Michael Schartner, Benjamin B. Ador, Francesca Simonelli, Acer Y.-C. Chang, Anil K. Seth

## Abstract

What are the global neuronal signatures of altered states of consciousness (ASC)? Recently, increases in neural signal diversity, compared to those found in wakeful rest, have been reported during psychedelic states. Neural signal diversity has previously been identified as a robust signature of the state of consciousness, showing lower scores during sleep or anaesthesia compared to wakeful rest. The increased neural signal diversity during psychedelic states raises the additional possibility that it may also reflect the increased diversity of subjective experiences associated with these states. However, psychedelic states involve widespread neuropsychopharmacological changes, only some of which may be associated with altered phenomenology. Therefore, we used stroboscopic stimulation to induce non-pharmacological altered states of consciousness while measuring the diversity of EEG signals. Stroboscopic stimulation caused substantial increases in the intensity and range of subjective experiences, with reports of both simple and complex visual hallucinations. These experiences were accompanied by increases in EEG signal diversity scores (measured using Lempel-Ziv complexity) exceeding those associated with wakeful rest, in line with studies of the psychedelic state. Our findings support the proposal that EEG signal diversity reflects the diversity of subjective experience that is associated with different states of consciousness.

## Introduction

The search for neural markers of consciousness has often been described as having two aspects. One relates to the state of consciousness, or conscious ‘level’, as in the contrast between the normal waking state and sleep or anaesthesia. The other relates to specific perceptual experiences, such as hearing a trumpet or smelling a rose, referred to as the ‘contents of consciousness’. Following early suggestions that conscious level may be related to the phenomenological diversity of experience^1, 2^, there has been increasing interest in characterising the interaction between conscious level and conscious content^3^, with recent attempts describing states of consciousness as regions within a multidimensional space^3, 4^.

Bayne et al. (2016) propose that one dimension along which conscious states differ is how they ‘gate’ conscious content. For example, the normal waking state allows for a wide range of contents to enter conscious experience, allowing for many unique cognitive and behavioural responses. However, in other states, such as mild sedation or during seizures, experiential content appears to be gated more restrictively, compared to the normal waking state, reducing the repertoire of cognitive and behavioural responses. The reduction in the diversity of subjective experiences during these ‘diminished’ states may plausibly be reflected in the diversity of neural signals in these states. Indeed, measures of spontaneous EEG signal diversity, such as Lempel-Ziv complexity, are reduced in states with diminished or absent conscious experience, such as during sleep or under anaesthesia, relative to the normal waking state ^5-7^. In a more direct test of the relation between experiential diversity and neuronal signal diversity, Schartner et al., ^8^ reported a brain-wide robust increase in signal diversity during altered states of consciousness (ASC) induced by the classic psychedelics LSD, ketamine and psilocybin. Notably, psychedelic drugs produce powerful phenomenological effects on an individual’s experience of self and world, with subjective reports suggesting that the psychedelic state is associated with a general ‘broadening’ of the range of visual perceptual contents (i.e. simple imagery, complex imagery and audio-visual synaesthesia) compared to the normal waking state ^9-11^. The results of ^8^ therefore demonstrate that neuronal signal diversity is not only sensitive to diminished or unconscious states, but also to the alterations in experiential content associated with psychedelic ASC, raising the interesting possibility that spontaneous signal diversity may show sensitivity to the diversity of subjective experience in general.

In addition to their powerful phenomenological effects, psychedelic drugs also induce a plethora of general physiological effects that may potentially contribute to alterations in signal diversity, such as increases in heart rate, plasma cortisol, oxytocin, epinephrine, and the like^12^. Psychedelics also have been shown to induce specific changes to the spectral properties of EEG, such as a pronounced decrease in alpha power (8-13 Hz)^13-15^. The extent to which these various factors are connected to altered perceptual phenomenology remains unknown^10, 13, 16, 17^. While alterations of the spectral properties of EEG can to some extent be controlled for by surrogate data methods^8^, it is harder to exclude the potential effects of an altered pharmacological milieu on neuronal signal diversity. Therefore, in the present study, we used stroboscopic stimulation to induce non-pharmacological ASC, while measuring the spontaneous diversity of EEG signals using single channel Lempel-Ziv complexity (LZs).

Stroboscopically-induced visual hallucinations (SIVH) were first reported by Jan Purkinje in 1819, who saw crosses, stars and spirals, when waving his hand between his eyes and the sun (Purkinje, 1819). SIVH occur with eyes closed and are characterised by intense visual experiences of amorphous colours, geometric patterns and colourful kaleidoscopic imagery^18-23^. The specific form of SIVH appears to depend on the stroboscopic frequency, for example, stroboscopic stimulation at <10 Hz tends to induce radial patterns, whereas stimulation between 10–20 Hz tends to induce spiral patterns^18, 24^. The most vivid visual experiences are usually reported to occur at a stimulation frequency close to the dominant alpha frequency (8-12 Hz), with stimulation frequencies < 5 Hz or > 30 Hz inducing subjectively weaker visual experiences^18, 24-26^. The contents of SIVH remain outside of voluntary control, but the individual usually retains insight into their non-veridical nature^25^. The same applies to some psychogenic or neurological ASC, such as those occurring in Charles Bonnet Syndrome^25, 27^. Additionally, as noted by several authors^22, 28-30^, SIVH are similar in aspects of perceptual content to the altered visual experiences induced by hallucinogenic drugs. Beyond these perceptual effects, stroboscopic stimulation has also been shown to induce a range of dissociative phenomena including temporary disruptions in awareness, memory or the sense of identity or self^31, 32^. These types of dissociative experiences have also been reported in the psychedelic state^33-37^.

In the present study, we first characterised the subjective effects of stroboscopic stimulation at different frequencies, using measures of the subjective intensity of experience and an adapted Altered States of Consciousness Questionnaire (ASCQ, ^14, 38^), a well validated and widely used tool for defining different ASC, such as psychedelic states. Next, to examine the similarities between psychedelic and stroboscopic ASC, we compared ASCQ responses following SIVH to those found following the administration of psilocybin (using data from a separate study). Finally, to test the hypothesis that EEG signal diversity correlates with increases in the diversity of experiences, we examined the effects of stroboscopic stimulation at two different frequencies on signal diversity and compared them to a baseline of wakeful rest (eyes closed with no stimulation). Based on the increases in neuronal signal diversity observed in the psychedelic state, we hypothesised that we would also see increases due to stroboscopic stimulation, if such stimulation gave rise to hallucination-like visual phenomena. We performed additional exploratory correlation analyses, to verify if LZs and each normalised spectral power band were reflecting distinct features of the data.

## Methods

### Participants

Twenty-three healthy students of the University of Sussex took part in this experiment (mean age of 23.4, seven male). Participants provided informed consent before taking part and received £10 or course credits as compensation for their time. To reduce the risks associated with stroboscopic light stimulation, participants were required to complete questionnaires on anxiety and epilepsy before taking part and were excluded if they met predefined exclusion thresholds for each (no participants met these exclusion criteria). Participants 01 and 12 were excluded due to missing data. Participants 19 and 20 were excluded due to excessively noisy EEG data, leaving 19 participants for all further analyses. The experiment was approved by and run in accordance with the University of Sussex ethics committee.

### Experimental design and data acquisition

Participants were seated in a dark electromagnetically shielded room 50 cm away from a Lucia N^0^03 stroboscope (Innsbruck, Austria, see Fig 1). Stroboscopic light stimulation was presented at 80% of maximum LED output, resulting in a maximum luminous flux of approximately 5600 lm (lumen) over the participant eyes. There were three stimulation conditions: ‘Dark’ (no stimulation), ‘3 Hz’ (strobe at 3 Hz) and ‘10 Hz’ (strobe at 10 Hz). Conditions were always presented in this order in both the practice session and in the main experiment. These frequencies were chosen based on previous research^18, 21, 23^ and the results of a pilot study (see Supporting Information), as they reliably elicited powerful phenomenological effects. Participants were instructed to keep their eyes closed during all conditions.

**Figure 1.**
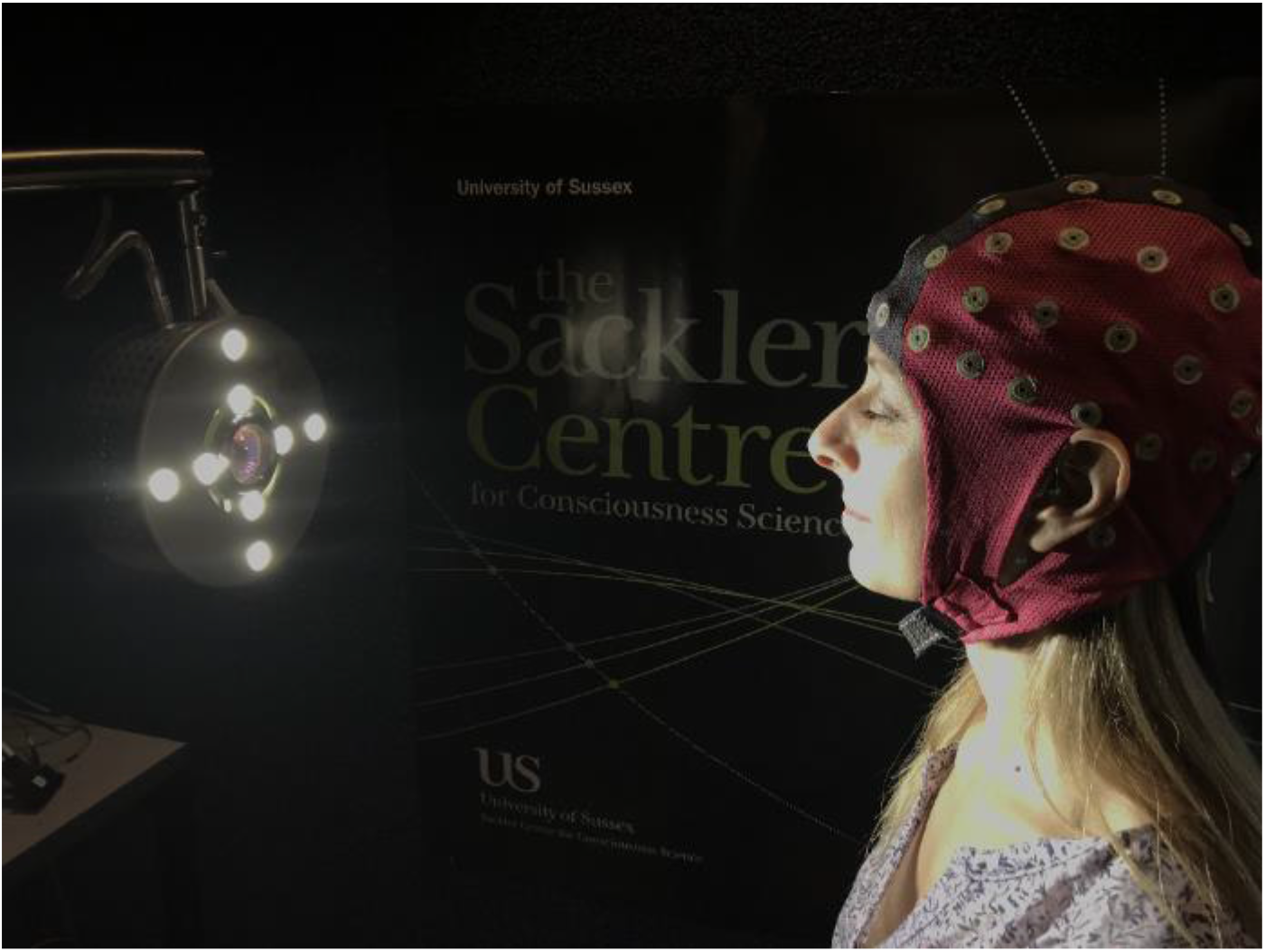
Lucia N^0^03 stroboscope used in the current experiment (Innsbruck, Austria), consisting of an array of 8 high power LEDs, and central halogen lamp (not used in this study), with a stroboscopic range of 0.004 Hz to 200 Hz. A maximum luminous flux of approximately 5600 lm (lumen) was used in this experiment. The participant is seated facing the stroboscope, 50 cm away, with eyes closed and wearing a 64 channel Waveguard EEG cap.

Before taking part in the main experiment, participants were given a practice session, in which they completed a brief 90 second demo of each of the three conditions (30 seconds per condition). During the practice, for each condition, participants were asked “Please rate the intensity of your experiences or conscious contents on a moment to moment basis from zero to five using the visual analogue slider”, using the following scale 0 = ‘Low Intensity’ to 5 = ‘High Intensity’. Data from the visual analogue slider (VAS) was recorded using custom scripts (MATLAB R2016b, Natick, MA). Participants were encouraged to adapt their ratings to use the full range of the scale to more accurately reflect their moment-to-moment experiences. The practice session was repeated if participants either reported that they were unsure how to respond or if they only used a small subset of the intensity scale across all three conditions. Following the practice session, participants completed a computer-based version of the Altered States of Consciousness Questionnaire (ASCQ, adapted from^14, 38^); participants were asked to base their responses on an average of their experiences across stroboscopic conditions in the practice session (See Fig 3).

The main experiment consisted of the same conditions ‘Dark’ (no stimulation), ‘3 Hz’ (strobe at 3 Hz) and ‘10 Hz’ (strobe at 10 Hz) presented sequentially in this order for all participants, each lasting for 10 minutes. Participants continuously rated the intensity of their conscious content using the VAS during all conditions. Following each condition, participants completed the ASCQ and were also asked separately to rate the intensity of their experience as an average over the ten minute session from 0 = ‘low intensity’ to 5 = ‘high intensity’. Additionally, participants were asked to verbally describe a summary of their visual experiences during each session, to verify if all participants experienced the ‘simple’ visual hallucinations (e.g. simple oriented lines, checkerboards, grids, spirals or honeycombs) commonly reported during stroboscopic stimulation. If participants reported that they had experienced ‘complex’ SIVH during a session, such as life-like objects or scenes, they were asked to describe their experiences in writing. They were then asked to rate the visual clarity of each ‘complex’ experience on a scale from one to ten, 1 = ‘not clear at all’, 10 = ‘as clear and vivid as veridical perception’. Completing these assessments meant that participants had a minimum of 5 min break (maximum 10 min) between each condition.

EEG data were recorded with ASA-Lab 4.7.11 using a 64 channel ANT Neuro amplifier at a sampling rate of 2048 Hz and a 64 channel Waveguard EEG cap (ANT Neuro, Enschede) employing standard Ag/AgCl electrodes placed according to the 10-20 system, using an average reference. No analogue filter was applied during online recording. Pre-processing of the EEG data was performed using the EEGLAB 12.0.2.6b toolbox (Delorme & Makeig, 2004) under MATLAB R2016b (Mathworks, Inc. Natick, MA, USA) and data analyses were performed using custom MATLAB, Python (www.python.org) and R 3.3.1 scripts. The acquired EEG data were down-sampled to 250 Hz and then filtered using a 1-30 Hz FIR bandpass filter. This frequency range was chosen due to the known susceptibility of LZs to high-frequency noise^39^. Artefact-contaminated channels were identified using EEGLAB based on the kurtosis of all recorded electrodes: electrodes with a z-score of >5 were removed and automatically interpolated using spherical interpolation of neighbouring electrodes. The assumptions for parametric statistics (normality or homoscedasticity) were tested on each data set to choose the correct statistical test.

### Analysis

#### Analyses ASCQ

To investigate the subjective effects of stroboscopic stimulation at 3 Hz and 10 Hz and to confirm that they elicited distinct phenomenological states we compared ASCQ responses following these two conditions. First, we averaged the absolute values (0-100) of all participants’ responses for each ASCQ dimension and condition. We used a repeated measures ANOVA consisting of factors ASCQ dimension and condition (Dark/3 Hz/10 Hz) to assess if there were differences in ASCQ responses between conditions. Following this primary analysis, we conducted a number of follow-up analyses to investigate between-condition differences across all ASCQ dimensions, as well as questionnaire and VAS measures of subjective intensity using frequentist and Bayesian paired *t*-tests. To explore potential phenomenal similarities between stroboscopic and psychedelic ASC, we compared the difference of average ASCQ dimensions for experimental and baseline conditions: 3 Hz-Dark, 10 Hz-Dark and psilocybin-placebo (psilocybin data taken from^14^) using frequentist *t*-tests Bonferroni corrected between conditions (denoted *p_bon_*), and Bayesian independent sample *t*-tests.

For all Bayesian tests we quantified proximity of the data to the null (no difference between results), or to the alternative hypothesis (difference in results), using JASP (JASP Team, 2018), with a default Cauchy prior of .707 half-width at half-maximum. A BF10 > 3.0 is widely interpreted as evidence for accepting the alternative hypothesis (i.e. there is a difference), whereas BF10 < 0.33 is widely interpreted as evidence for accepting the null hypothesis (i.e. there is no difference)^40^.

#### EEG signal analysis

Single channel Lempel-Ziv complexity (LZs) was used to assess the temporal EEG signal diversity of the pre-processed EEG data, following the implementation described in^8^. For a given segment of data, LZs quantifies the diversity of the signal by counting the number of distinct patterns of activity in the data (see Fig 2 and see https://github.com/mschart/SignalDiversity for python code to compute LZs from EEG data). Computing LZs requires the binarization of the multidimensional time series. Briefly, the signal diversity of each EEG channel is assessed independently per condition by dividing each 10 min recording into 10 s non-overlapping time series segments, subtracting the mean amplitude from each segment, dividing by standard deviation and then removing any linear trend using the scipy signal detrend function.

**Figure 2.**
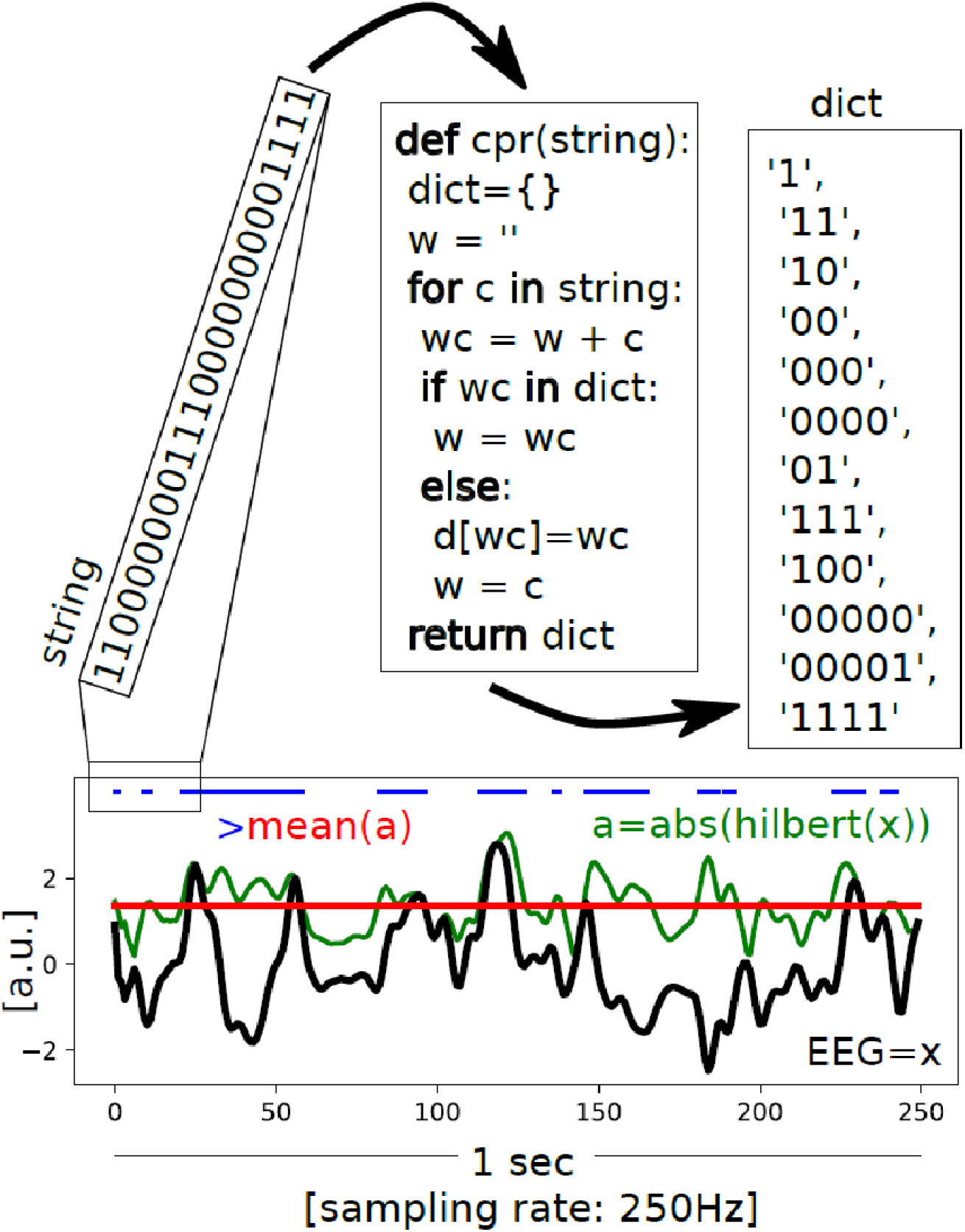
Schematic of the LZs computation. An example EEG signal with a sampling rate of 250 Hz and length of 1 sec is shown in black (x). The mean (red) of the absolute value of its analytic signal (green, a) is used to binarise the signal (blue). The encoding step of the Lempel-Ziv algorithm is then applied to the first 25 entries of that binarized signal (in this illustration), creating a dictionary of the unique subsequences, which is then normalized by dividing the raw value by those obtained for the same randomly shuffled binary sequence. This provides a value between 0-1 that quantifies the temporal diversity of the EEG signal (LZs).

**Figure 3:**
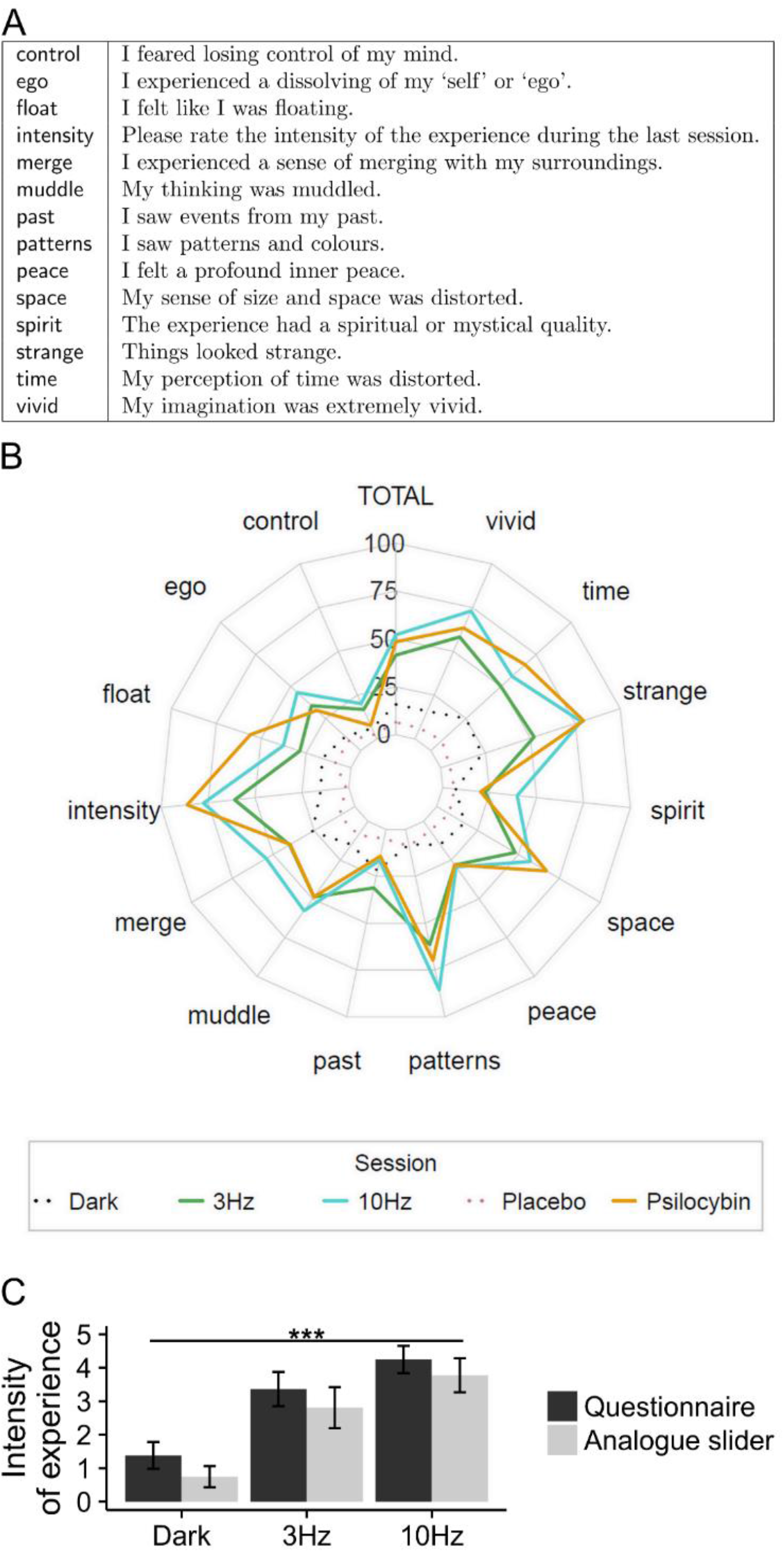
Summary of ASCQ responses. **A.** ASCQ dimensions and questions. **B.** Radar plot of ASCQ responses, averaged across participants. Responses for stroboscopic conditions, 3 Hz (‘green’), 10 Hz (‘blue’) and Dark condition (‘black dotted line’), along with psilocybin (‘orange’) and placebo (‘purple-dotted line’) responses (data for psilocybin and placebo taken from^14^). **C.** Subjective ratings of intensity of experience, averaged across participants, taken from absolute values of post-hoc reports (ASCQ, ‘black’) and ‘live’ visual analogue slider responses (‘grey’) obtained during each condition. The subjective intensity of experience for both stroboscopic conditions was substantially higher than for the Dark condition. Statistics show Kruskal-Wallis rank sum test on both questionnaire (*p* = 8.8 × 10^−8^) and average analogue slider responses (*p* = 9.0 × 10^−8^).

The continuous signal of each channel is then transformed into a string of binary digits, using the mean of its absolute value as a threshold (based on the amplitude of the analytic signal), resulting in a sequence of zeros and ones for each channel. The LZs algorithm divides the sequence into non-overlapping and unique binary subsequences. LZs complexity is proportional to the number of unique binary subsequences: the greater the degree of randomness, the greater the number of different subsequences that will be present, and thus the higher LZs. The LZs score for each segment is normalized by dividing the raw value by the value obtained for the same binary input sequence that is randomly shuffled. Since the value of LZs for a binary sequence of fixed length is maximal if the sequence is entirely random, the normalized values indicate signal diversity on a scale from 0 to 1. All plots reflect the average (normalised) LZs score across all EEG channels.

We next computed LZs for two surrogate data types to test the relative contributions of the power and phase spectra to signal diversity during SIVH. The first surrogate data were derived by applying a fast Fourier transform to the original data and then randomising the phases of the sinusoids, while keeping their amplitudes fixed, before transforming back using an inverse Fourier transform^41^. This procedure produces surrogate time series data which have a random phase spectra, while retaining the power spectra of the original data. Therefore, for this surrogate data, only cross-condition differences in power spectra remain, allowing us to quantify the contribution of the power spectrum on signal diversity scores across conditions. These surrogate data are denoted by ‘EqualPhase’.

For the second surrogate data set we implemented a novel approach, which equalized the power spectra across all experimental conditions as follows. For each 10 s segment and channel, the power spectrum of the Dark condition was obtained via a discrete fast Fourier transform and used to replace the amplitudes of the sinusoids for the corresponding time series segment for both stroboscopic conditions. This resulted in the power spectrum of any segment of data from the 3 Hz and 10 Hz conditions being identical to the corresponding segment from the Dark condition, while the phase spectra of all segments remained unchanged. Therefore, for this surrogate data, any observed differences in LZs between conditions must be due to between-condition differences in the phase spectra, allowing us to estimate the contribution of the phase-spectrum to signal diversity scores across conditions. These surrogate data are denoted by ‘EqualPower’.

We used a repeated measures ANOVA consisting of factors condition (Dark/3 Hz/10 Hz) and Data-type (original/EqualPhase/EqualPower) to compare differences in LZs. Following this primary analysis, we conducted a number of follow-up analyses to investigate between-condition differences in LZs, as well as difference in LZs between Data-type using frequentist paired t-tests, Bonferroni, corrected for multiple comparisons.

Each condition was presented for 10 min in total, raising the possibility that subjective experiences and electrophysiological signals may have changed over this timeframe. To investigate how signal diversity changed over time (during each condition) we conducted an exploratory analysis. We computed the temporal evolution of LZs by averaging LZs scores across all electrodes for each 60 second non-overlapping segments for each condition. We then computed a least-squares regression of LZs by time for each condition. To better control for the variance between participants, for all time plots within-subjects errors were calculated using the method described in^42^, in which standard errors are taken from normalized data. The data was normalized by subtracting from each single-subject value the mean for that subject across conditions and then adding the overall mean. Since standard errors of normalized data are smaller on average than non-normalized data, the normalized data was then adjusted using a correction factor. Error bars computed following this procedure better control for the variance between subjects.

##### Normalized spectral power

Further exploratory analyses were carried out to compare signal diversity measures with more standard electrophysiological measures. We computed the average spectral power profile for delta (1-4 Hz), theta (4-8 Hz), alpha (8-13 Hz) and beta (13-30 Hz) frequency bands, for each condition and participant. As for LZs, for each participant and condition the 10 min recording was divided into non-overlapping 10 s segments. For each 10 s segment and electrode the spectral density of each frequency band was computed using the discrete fast Fourier transform, (https://github.com/mschart/SignalDiversity). Data from each frequency band was normalised by the sum of the power of all 4 bands, resulting in each frequency band representing a percentage of the total spectral power (i.e., relative power, not absolute power). The data was then averaged across segments, electrodes and finally participants. We compared changes in power spectra for each frequency band across conditions using paired sample t-tests. In addition, we computed a time-frequency decomposition of the EEG signal for each condition, averaged across participants and electrodes, using the EEGLAB newtimef function. Spectral power (μV^2^) was converted to the dB scale using a log10 conversion, using a 1000ms time-window prior to the beginning of each 10 minute session as the baseline. We examined changes in each frequency band over time and condition, in the same manner as described for LZs. To test for significant alterations in each frequency band over time we computed a least-squares regression for each condition and frequency band.

##### Associations between signal diversity and power spectra

To test for relations between LZs and the power spectrum, for each condition we computed a Pearson correlation (r) across participants using an average of LZs scores and normalised spectral power values for delta, theta, alpha and beta frequency bands. Correction for multiple comparisons was controlled for using Bonferroni correction.

## Results

### Altered States of Consciousness Questionnaire (ASCQ) and intensity ratings during SIVH

The dimensions and responses to the ASCQ for each condition are shown in Fig 3. A two-factorial repeated measures ANOVA consisting of factors ASCQ dimension and condition (Dark/3 Hz/10 Hz) revealed a significant interaction between these factors (F(18,1) = 6.171, *p* < .001, η^2^ = 0.153) and a significant difference between ASCQ dimensions (F(18,1) = 38.44, *p* < .001, η^2^ = 0.318). Post-hoc paired *t*-tests Bonferroni corrected for multiple comparisons revealed a significant difference in ASCQ responses between Dark and 3 Hz (t(18)= 4.09, *p_bon_* < .001) and Dark and 10 Hz (t(18)= 5.93, *p_bon_* < .001). There were no significant differences in ASCQ responses between 3 Hz and 10 Hz (t(18)= 1.84, *p_bon_* = 0.42) with both conditions being highly correlated (r = 0.95, *p* = 1.5 × 10^−7^), indicating some commonality in subjective experiences between these conditions. Next, we examined the differences in individual ASCQ dimensions between stroboscopic conditions, finding that 3 Hz and 10 Hz differed significantly on the following dimensions: ‘Ego’, ‘Float’, ‘Intensity’, ‘Merge’, ‘Patterns’, ‘Space’, ‘Spirit’, ‘Strange’ and ‘Vivid’ (see Table 1). To quantify these differences, we conducted additional Bayesian paired *t*-tests. The analysis provided further evidence in support of the alternative hypothesis (BF10 > 3) i. e. a difference in ASCQ scores for 3 Hz and 10 Hz for the same dimensions as found above (see Table 1). Across all ASCQ dimensions ‘Peace’ was the only dimension to show evidence in support of the null hypothesis between 3 Hz and 10 Hz.

**Table 1.** Bayesian and standard statistical comparisons of ASCQ ratings between 3 Hz and 10 Hz conditions (paired *t*-tests, Bonferroni corrected), 10 Hz and psilocybin and 3 Hz and psilocybin (independent *t*-tests, Bonferroni corrected). Dagger symbols and bold text indicates Bayes Factor values in excess of 3, which show evidence in favour of a difference between ASCQ responses. Asterisks after *p*-value indicates a significant difference in standard Bonferroni corrected *t*-tests. For comparisons between studies, statistics were applied on the difference of experimental and baseline conditions: 3 Hz-Dark, 10 Hz-Dark and psilocybin-placebo.

| ASCQ<br>Dimension | 10 Hz vs<br>3 Hz |  | 10 Hz vs<br>Psilocybin |  | 3 Hz vs<br>Psilocybin |  |
| --- | --- | --- | --- | --- | --- | --- |
|  | BF<sub>10</sub> | *p*<sub>bon-value</sub> | BF<sub>10</sub> | *p*<sub>bon-value</sub> | BF<sub>10</sub> | *p*<sub>bon-value</sub> |
| Control | 0.331 | 0.75 | 0.48 | 0.61 | 0.387 | 0.5 |
| Ego | **3.34†** | 0.017* | 0.32 | 0.97 | 0.429 | 0.43 |
| Float | **8.09†** | 0.006* | **5.35†** | 0.009* | **57.24†** | 8.6 × 10<sup>-4*</sup> |
| Intensity | **56170†** | 2.8 × 10<sup>-7*</sup> | **7.63†** | 0.006* | **970†** | 4.4 × 10<sup>-5*</sup> |
| Merge | **3.96†** | 0.014* | 0.46 | 0.29 | 1.462 | 0.041 |
| Muddle | 0.58 | 0.17 | 0.36 | 0.64 | 0.608 | 0.22 |
| Past | 0.84 | 0.1 | 1.43 | 0.06 | 0.408 | 0.55 |
| Patterns | **1130†** | 2.1 × 10<sup>-5*</sup> | 0.40 | 0.79 | 0.836 | 0.083 |
| Peace | 0.25 | 0.76 | 0.55 | 0.25 | 0.558 | 0.068 |
| Space | **4.50†** | 0.009* | **3.57†** | 0.016* | **32.85†** | 0.009* |
| Spirit | **6.75†** | 0.008* | 0.475 | 0.34 | 0.44 | 0.4 |
| Strange | **73.27†** | 4.6 x 10<sup>-4</sup>* | 0.459 | 0.36 | **142†** | 1.2x10<sup>-4</sup>* |
| Time | 0.62 | 0.16 | **3.88†** | 0.014* | **12.16†** | 0.0004* |
| Vivid | **5.90†** | 0.009* | 0.96 | 0.105 | **20.01†** | 0.002* |
| TOTAL | **219.66†** | 0.0001* | 1.12 | 0.098 | **22.13†** | 0.002* |

Examining the intensity of experience during each condition using questionnaire scores, we found that participants rated 3 Hz as being more intense than Dark (Wilcoxon rank sum test with continuity correction, *p* = 1.715 × 10^−5^), and 10 Hz rated as substantially more intense than the 3 Hz (*p_bon_* = .006). The intensity ratings scores given moment-to-moment during the experiment, using the analogue slider, averaged across the 10 min time window, revealed a similar pattern of results, with higher intensity ratings during 3 Hz (*p_bon_* < .001) and 10 Hz (*p_bon_* < .001) compared to Dark, as well as for 10 Hz compared to 3 Hz (*p_bon_* < .001). Averaging across all ASCQ dimensions as a measure of overall intensity^41^, ‘TOTAL’, revealed a significant difference between Dark and 3 Hz (t(18)= 4.19, *p_bon_* < .001) and Dark and 10 Hz (t(18)= 6.21, *p_bon_* < .001), and also between 3 Hz and 10 Hz, (t(18)= 1.81, *p_bon_* < .001). Participants found 10 Hz more intense than 3 Hz, with both stroboscopic conditions being more intense than the Dark condition.

Together, these results indicate that stroboscopic stimulation at 3 Hz and 10 Hz led to marked increases in the intensity and diversity of subjective experiences compared to Dark. While there were some commonalities in experience between 3 Hz and 10 Hz, they also differed in terms of intensity and across many ASCQ dimensions, indicating that each stimulation frequency produced distinct phenomenal states.

We next compared ASCQ responses for 3 Hz and 10 Hz to those given following the administration of psilocybin (data taken from^14^, Fig 3B and Table 1). To allow an interpretable comparison between the results of our study and previous results^14^, all analyses used the difference of each ASCQ dimension for experimental and baseline conditions, i.e. 3 Hz-Dark, 10 Hz-Dark and psilocybin-placebo. The phenomenological profile of both 3 Hz and 10 Hz displayed marked similarities across multiple phenomenal dimensions to the psilocybin-induced psychedelic state (Figure 3B; 3 Hz: r = 0.87, *p* = 5.6 × 10^−5^; 10 Hz: r = 0.83, *p* = 2.1 × 10^−4^; Pearson’s product-moment correlation, Bonferroni corrected). While there was a significant difference in the post-hoc reported intensity experience during both 3 Hz and 10 Hz compared to psilocybin (Table 1), we found no significant difference in responses for ‘Control’, ‘Ego’, ‘Merge’, ‘Muddle’, ‘Past’, ‘Patterns’, ‘Peace’, ‘Spirit’ ASCQ dimensions (Table 1). Additionally, 10 Hz did not differ significantly from psilocybin responses for the ‘Vivid’ and ‘Strange’ ASCQ dimensions. Overall intensity, ‘TOTAL’ (average of all ASCQ dimensions), was higher for psilocybin than 3 Hz (*p* = 9.2 × 10^−4^, Welch Two sample *t*-test). However, 10 Hz and psilocybin did not differ significantly on this measure (*p* = 0.098). We next quantified how close to the null hypothesis 10 Hz and psilocybin ASCQ responses were using independent Bayesian *t*-tests, which revealed evidence in support of the null hypothesis (i.e., no difference) only for the ‘Ego’ dimension. The analysis provided evidence in support of the alternative hypothesis for the following dimensions: ‘Float’, ‘Intensity’, ‘Space’, ‘Time’. For the remaining dimensions, Bayesian analyses were insensitive as to whether the null or alternative hypothesis was supported (see Table 1). Performing the same analysis for 3 Hz and psilocybin, we found evidence in support of the alternative hypothesis for the following ASCQ dimensions: ‘Float’, ‘Intensity’, ‘Space’, ‘Strange’, ‘Time’, ‘Vivid’ (Table 1). Together, these results suggest that the changes in experience and phenomenal content – as reflected by the ASCQ – reported during 3 Hz and 10 Hz stroboscopic stimulation showed some similarities to those following the ingestion of psilocybin, but also a number of differences. These results support stroboscopic stimulation as a novel non-pharmacological method of inducing ASC, phenomenologically similar in some respects to the psychedelic state. However, we acknowledge that post-hoc questionnaires are imperfect tools when attempting to capture a detailed description of phenomenal experience.

### Stroboscopically induced visual hallucinations

All participants verbally reported the occurrence of SIVH during both 3 Hz and 10 Hz stimulation, including reports of geometric patterns, spiral, grids and colourful kaleidoscopic imagery. The descriptions of these phenomena are consistent with previous reports of SIVH^18, 22, 23, 31, 43^. As mentioned, SIVH were reported as being more intense during 10 Hz compared to 3 Hz. Strikingly, in addition to these simple visual phenomena, the majority of participants (12 out of 19) also reported complex visual hallucinations (CVH), such as realistic scenes or faces^25, 43^. Participants described these experiences as building throughout the session, with the majority occurring toward the end of the 10 min period. 11 out of 12 reported that these experiences occurred during 3 Hz, with only 3 participants reporting that they also occurred during 10 Hz, despite the increase in ‘intensity’ reported in this condition. The remaining participant reported that CVH *only* occurred during 10 Hz. We reproduce below representative post-hoc reports of 3 Hz CVH, along with ratings of their visual clarity (relative to veridical perception, scale from 1-10, 1 = ‘not clear at all’, 10 = ‘as clear as veridical perception’):

**Participant 10.** “Viewing from below, I saw a body falling into water (reminded them of the intro to the Vikings TV Series). I saw a spaceship coming through a wormhole (centre of Lucia) – reminded me of Stargate. I saw House MD looking at me from above, he was curious”, 8/10.
**Participant 12.** “I saw historical faces. I saw an MRI scan projected in front of me. I saw a cityscape of London - with Big Ben”, 8/10.
**Participant 17.** “I had very vivid hallucinations of faces - friends, family, but also strangers – I could make out their individual features - blond hair, what they were wearing etc.”, 7/10.
**Participant 22.** “I was standing in a parking lot - although I couldn’t see any distinctive cars - with two gentlemen walking towards me, they were dressed identically - wearing a white top and brown trousers”, 10/10.

Two distinct types of CVH were reported. The first occurred only during 3 Hz and was comprised of detailed perceptions of faces, objects or scenes, which were described as being subjectively close to veridical perception, scoring high on visual clarity (M = 7.1 *SE* = 0.32, described above). The second, which more commonly occurred during 10 Hz, lacked the high-level of detail associated with 3 Hz CVH: “the impression of being in a graveyard”, “a hospital corridor”, “a spaceship view, going through hyperspace”, “flying through space in the chair”. These hallucinations were described as being distinct from veridical perception, with a correspondingly low visual clarity score (*M* = 4.3, *SE* = 0.33). While this second type of hallucination appears to be distinct from the ‘simple’ geometric hallucinations elicited by stroboscopic stimulation, the descriptions of these experiences do not meet the clinical criteria necessary to be classified as CVH, a term reserved for visual hallucinations of more complex phenomena such as figures and faces^25, 43^.

### Changes in EEG signal diversity during stroboscopic stimulation

In order to test our main hypothesis that neuronal signal diversity is sensitive to changes in phenomenological diversity, we compared EEG Lempel-Ziv scores (LZs) (schematically shown in Fig 2) using a repeated measure ANOVA consisting of main factors Condition (Dark/3 Hz/10 Hz) and Data-type (Original/EqualPhase/EqualPower). Results of the ANOVA revealed a significant main effect of Condition (F(2,36) = 36.9, *p* < .001, η^2^ = 0.672) and Data type (F(2,36) = 77.26, *p* < .001, η^2^ = 0.811) and a significant interaction between these factors (F(4,72) = 5.47, *p* < .001, η^2^ = 0.233). Post-hoc paired *t*-tests Bonferroni corrected for multiple comparisons revealed a significant increase in LZs at the group level for both 3 Hz (*t*(18)= −5.56, *p_bon_* = < .001) and 10 Hz (*t*(18)= −8.76, *p_bon_* = < .001) relative to Dark (Fig 4). However, comparing LZs scores between 10 Hz and 3 Hz we found that the difference did not survive correction for multiple comparisons (*t*(18)= −2.26, *p* = .01, *t*(18)= −2.26, *p_bon_* = .11) (see Fig 4). We found that signal diversity results were also robust at the single participant level (see Fig S7). For topographic distribution of between-condition changes in LZs see Fig S2. Together these results suggest that EEG signal diversity reflects the changes in brain dynamics associated with alterations in phenomenological diversity.

**Figure 4.**
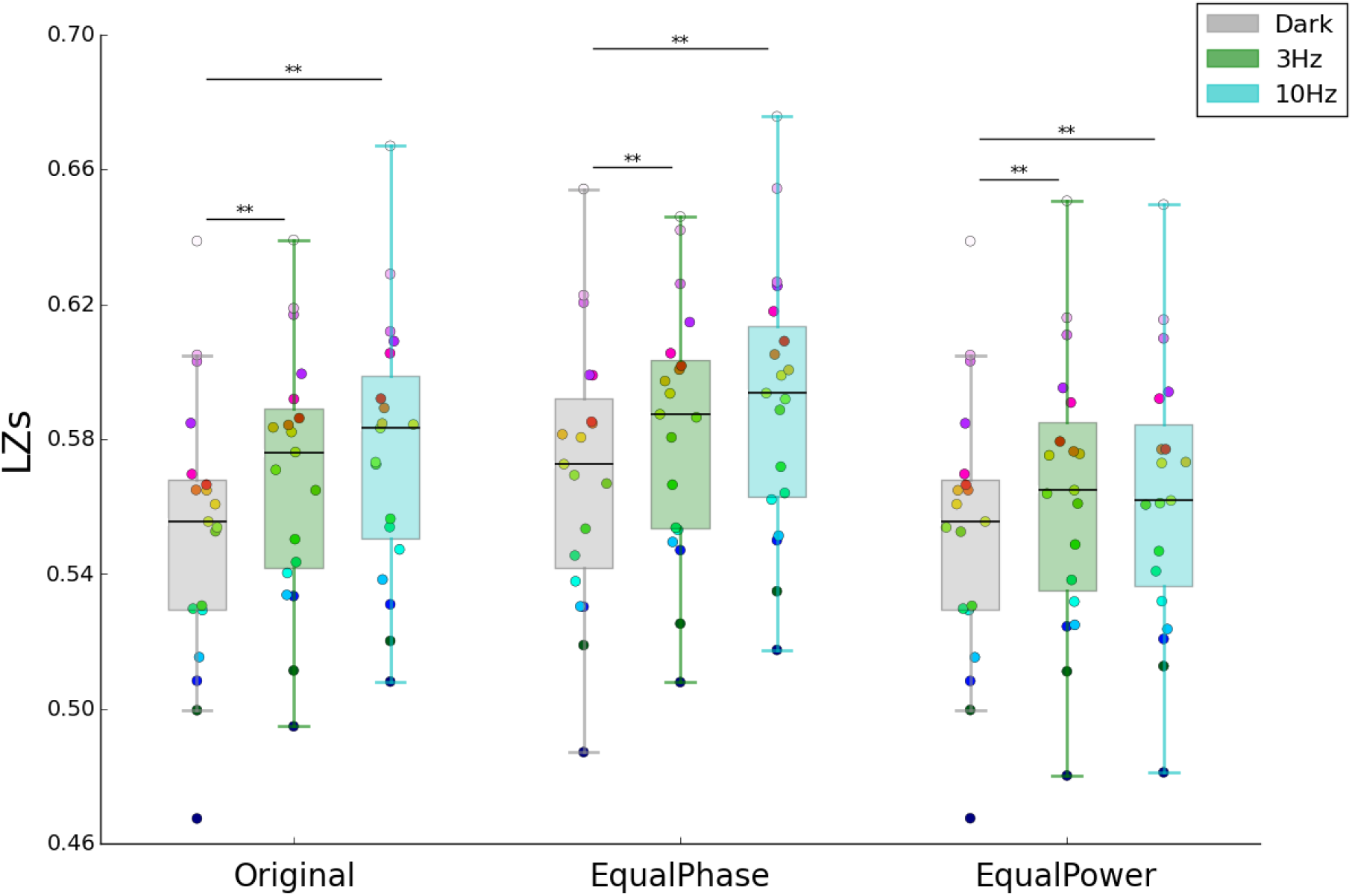
Increased EEG signal diversity during SIVH. Original: mean signal diversity for Dark, 3 Hz and 10 Hz. EqualPhase: mean signal diversity for surrogate phase-randomised data. Note, LZs scores are higher for all conditions, since phase randomisation by construction increases signal diversity. EqualPower: mean signal diversity for surrogate data with power-spectra equalised across conditions to Dark power-spectrum. The boxplots range extends from the lower to upper quartile values of the data, with a horizontal line at the median. Whiskers display the range of the data (excluding outliers). Coloured circles show individual participant results, with each unique colour representing the same participant across conditions for all data types. ** indicates a *p*-value < 0.01, obtained using paired sample *t*-tests, Bonferroni corrected for multiple comparisons.

Next, to determine the contribution of the power spectra and, separately, phase spectra to changes in signal diversity we performed post-hoc comparisons between LZs scores for Data type, (original, EqualPhase, testing for the effect of power and EqualPower, testing for the effect of phase). These tests revealed a significant difference in LZs scores between original and EqualPower (*t*(18)= −2.88, *p_bon_* = .003), original and EqualPhase (*t*(18)= −20.76, *p_bon_* < .001) and EqualPhase and EqualPower data (*t*(18)= −10.29, *p_bon_* < .001).

Examining changes in LZs for each surrogate dataset we found that EqualPhase data produced a similar pattern of signal diversity scores to the original data (Fig 4): a significant increase for both 3 Hz and 10 Hz compared to Dark (3 Hz, *t*(18) = −3.53, *p_bon_* = .007 and 10 Hz *t*(18) = −5.97, *p_bon_* < .001). 10 Hz and 3 Hz did not differ significantly (*t*(18) = −2.48, *p_bon_* = .07). Similarities in the pattern of LZs scores for each condition between the original and EqualPhase data (Fig. 4) indicates that between-condition changes in signal diversity can be attributed in part to changes in power spectrum. Note that the general increase in LZs for EqualPhase, across all conditions, is due to randomisation of the phase-spectra, which has the effect of increasing signal diversity by construction.

For EqualPower surrogate data, we found higher LZs scores for both 3 Hz and 10 Hz compared to Dark (Fig 4), (3 Hz, *t*(18)= −8.98, *p_bon_* < .001 and 10 Hz, *t*(18)= −7.53, *p_bon_* < .001), demonstrating that increases in EEG signal diversity in these conditions can be attributed in part to changes in phase spectra (since power spectra are equalized for these surrogate data). LZs scores for EqualPower were not significantly different between 3 Hz and 10 Hz (*t*(18) =1.92, *p_bon_* = .21).

EEG responses recorded over visual cortex have been shown to display a sharply peaked responses at the same frequency as the driving frequency, known as the steady state visual evoked potential (SSVEP)^21, 23^. To exclude the possibility that SSVEP could be responsible for the between-condition alterations in LZs, in a follow-up analysis we applied periodic Butterworth notch filters (width = 0.5Hz, order = 4 or 5 (3 Hz and 10 Hz, respectively) at the appropriate stroboscopic stimulation frequencies (3 Hz or 10 Hz) prior to computing LZs (Fig S5). Similar to the results of the original data, we found increases in signal diversity for both 3 Hz (Notch 3 Hz: t(18) = −6.02, *p_bon_* < .0001; Notch 10 Hz: t(18) = −3.72, *p_bon_* < .0001) and 10 Hz notch-filtered data (Notch 3 Hz: t(18) = −6.79, *p_bon_* < .0001; Notch 10 Hz: t(18) = −4.45, *p* < .001) relative to Dark. There was no significant difference in LZs between 3 Hz and 10 Hz for the notch-filtered data (Fig S5).

Besides its application to different sleep stages^7^, previous research has not explored the variation in LZs over extended periods of time, such as 10 min. To better understand how alterations in experience over time affect signal diversity, we investigated, in an exploratory analysis, the temporal evolution of LZs throughout each session by comparing LZs scores across non-overlapping 60-second windows, for each condition and data type (Fig 5). Using the original data, a least-squares regression of LZs scores by time revealed a significantly increasing linear trend in signal diversity for Dark (*r* = 0.94, *p* = < .001). However, in contrast to our main result of increased signal diversity scores for 3 Hz and 10 Hz conditions relative to baseline (Fig 4), we found a strong decreasing linear trend in LZs over the course of the experiment for both 3 Hz and 10 Hz (*r* = 0.86, *p* = < .001 and *r* = 0.68, *p* < .05 respectively, Fig 5).

**Figure 5.**
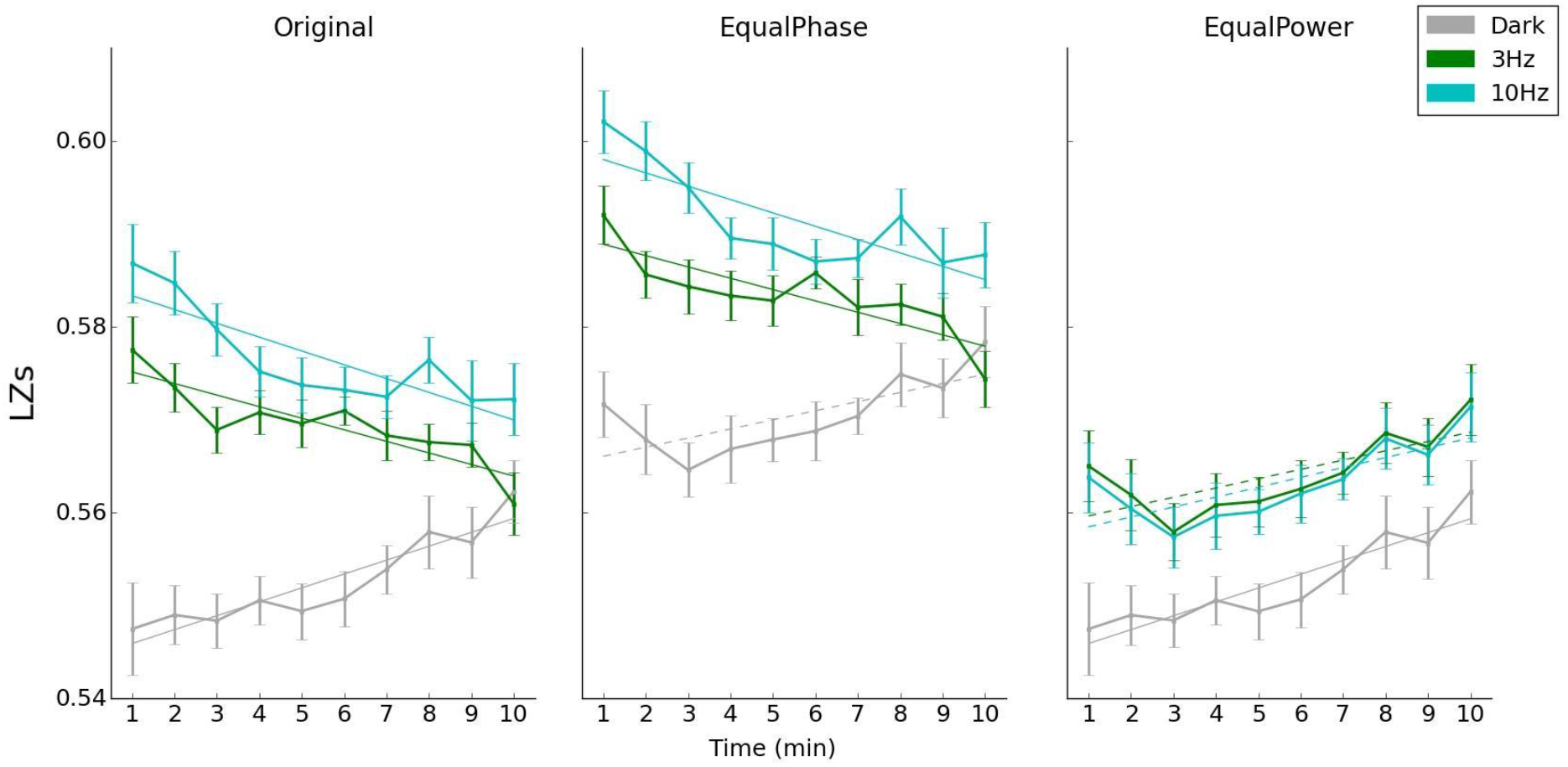
Mean signal diversity (LZs) across time for the original and two surrogate data sets (EqualPhase, EqualPower). Each point represents a non-overlapping 1-minute segment of data, across a total of 10 minutes. Dashed lines indicate least-squares regression *p*-values between 0.01 and 0.05, solid lines indicate *p* < 0.01. Error bars are normalised within-subjects^42^.

We repeated the same analysis on our two surrogate data sets. EqualPhase displayed a similar pattern, with LZs increasing over time for Dark (*r* = 0.72, *p* = < 0.01, Figure 5), and decreasing for both 3 Hz (*r* = 0.81, *p* = .004) and 10 Hz (*r* = 0.66, *p* = .04). In contrast, for all 3 conditions EqualPower showed an increase in LZs with time (Dark, *r* = 0.94, *p* < .0001; 3 Hz, *r* = 0.74, *p* = .02; 10 Hz *r* = 0.77, *p* < .01), again suggesting a strong influence of power spectrum on signal diversity. However, note that the temporal evolution of the LZs scores are not identical across conditions for EqualPower, which must be due to between-condition differences in the phase spectra.

Finally, to directly compare how alterations in experience affect signal diversity, we plotted the temporal evolution of both LZs and subjective ratings of the intensity of experience given in real-time using a visual analogue slider (VAS) (Fig S8). We found that for all conditions, on average, intensity increased throughout the session with no obvious relationship between these two measures across the 10 minute time window.

Summarising so far, both 3 Hz and 10 Hz stroboscopic stimulation caused signal diversity scores exceeding those associated with wakeful rest (Dark), a similar pattern of results as found for pharmacologically-induced psychedelic states^8^. These results further support the sensitivity of neural signal diversity to changes in brain dynamics associated with alterations in the diversity of experience. Using surrogate data, we were able to identify the relative contributions of the power and phase spectrum to LZs, showing that both factors contributed to increases in LZs (for 3 Hz and 10 Hz stimulation). Finally, when investigating changes in signal diversity over time using original data, surprisingly we found a clear decreasing trend of LZs for 3 Hz and 10 Hz, while, conversely, LZs scores increased with time during the Dark condition.

### Stroboscopic-induced changes in spectral power profile

To investigate the effects of stroboscopic stimulation on spectral power we computed a time-frequency decomposition of the EEG signal for each condition (Fig 6) . In line with previous reports, the stimulation frequency was strongly visible in the EEG, i.e. we observed the effects of entrainment of the EEG signal by stroboscopic stimulation at 3 Hz and 10 Hz, along with their accompanying harmonics (Figs 6 and S4)^18, 21, 26^.

**Figure 6.**
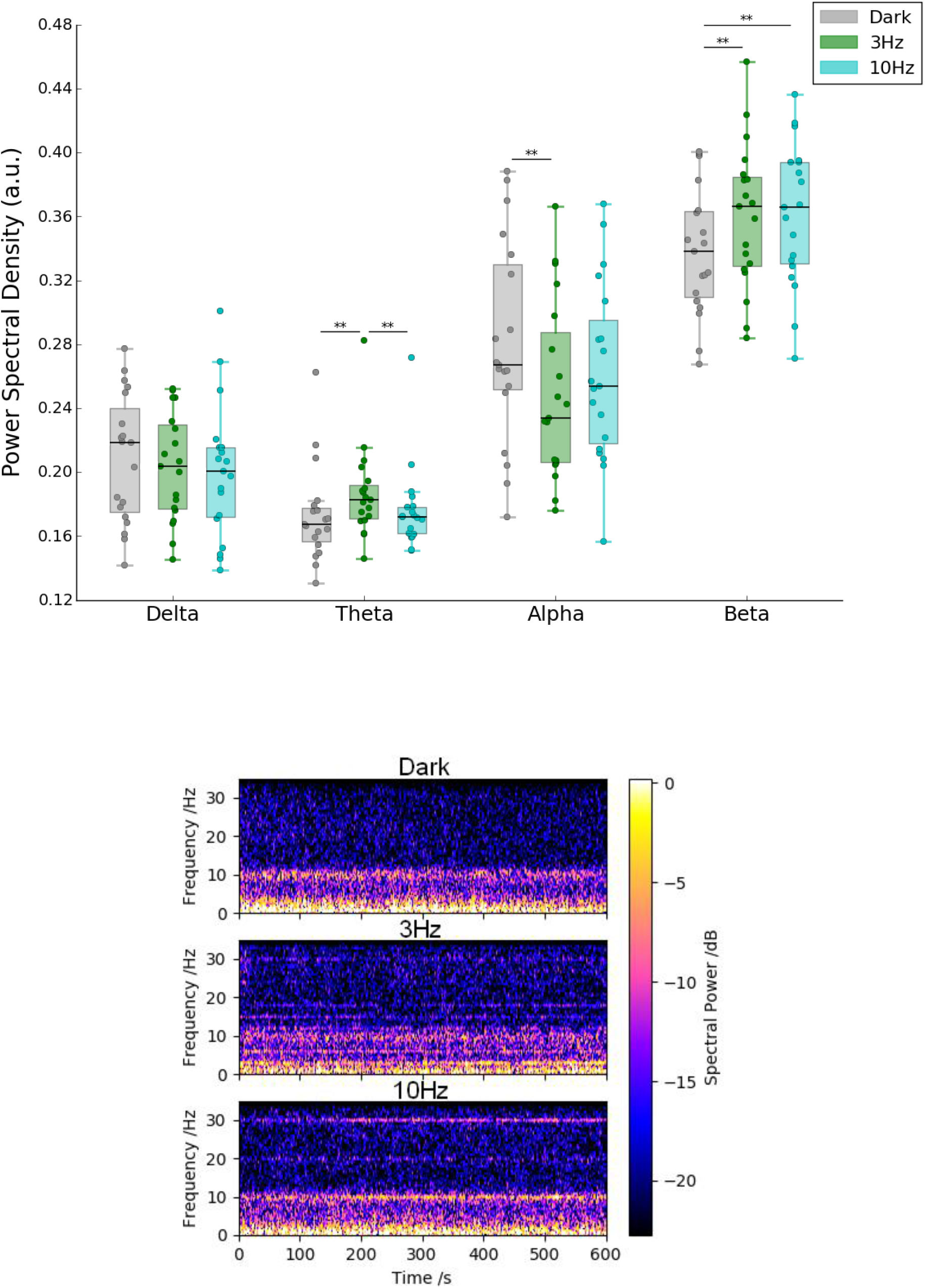
The top panel displays the average power spectral density for each condition and frequency band: delta (1-4 Hz), theta (4-8 Hz), alpha (8-13 Hz) and beta (13-30 Hz). The box extends from the lower to upper quartile values of the data, with a horizontal line at the median. Whiskers display the range of the data (excluding outliers, represented by diamond-shaped symbols). ** indicates a *p*-value < 0.01, obtained using paired sample *t*-tests, Bonferroni corrected. Units are power spectral density, measured in arbitrary units (a.u.), normalised by the sum of the spectral power of all frequency bands, resulting in each frequency band representing a percentage of the total spectral power. The bottom panel displays a time-frequency decomposition of the EEG signal for each condition averaged across participants and electrodes. Note the entrainment of the EEG signal due to stroboscopic stimulation at 3 Hz and 10 Hz and the accompanying harmonic frequencies, leading to an SSVEP. For an analysis of the effects of SSVPs on LZs see Fig S4 and S5. Spectral power (μV^2^) was converted to the dB scale using a log10 conversion. Topographic plots displaying the difference in average power spectral density for each frequency band and condition are displayed in Fig S3.

Next, we computed the average spectral power profile, using original data, for the delta (1-4 Hz), theta (4-8 Hz), alpha (8-13 Hz) and beta (13-30 Hz) frequency bands for each condition. As shown in Fig 6, we observed a substantial decrease in alpha power during 3 Hz (t(18) = 3.76, *p_bon_* = . 0043), but not 10 Hz (t(18) = 2.29, *p_bon_* = .010) relative to Dark. We note that a marked decrease in alpha power has previously been reported for psychedelic ASC^8, 10, 13, 14^. Beta power increased during both 3 Hz (t(18) = −5.37, *p_bon_* < .001) and 10 Hz (t(18) = −7.42, *p_bon_* < .001) relative to Dark. Finally, theta power increased during 3 Hz compared to both Dark (t(18) = −4.81, *p_bon_* < .001) and 10 Hz (t(18) = 3.71, *p_bon_* = .004). We observed no significant differences in average delta power across conditions.

Next we analysed the temporal evolution of the spectral power profile for the delta (1-4 Hz), theta (4-8 Hz), alpha (8-13 Hz) and beta (13-30 Hz) frequency bands for each condition, by considering non-overlapping 60-second segments of data (Fig 7). A least-squares regression of the spectral density of each frequency band by time revealed a significantly increasing linear trend in delta for 3 Hz (*r* = 0.69, *p* = .03). All conditions showed an increasing trend in theta, with 3 Hz showing the largest increase (Dark (*r* = 0.98, *p* < .0001), 3 Hz (*r* = 0.95, *p* < .0001), 10 Hz (*r* = 0.87, *p* = .001). We observed a decreasing linear trend for alpha only for 3 Hz (*r* = −0.77, *p* = .01) and Dark (*r* = −0.83, *p* = .003). Finally, there was a significant decreasing linear trend in beta spectral density for 3 Hz (*r* = −0.88, *p* = .0009), and an increasing trend for Dark (*r* = 0.83, *p* = .003).

**Figure 7.**
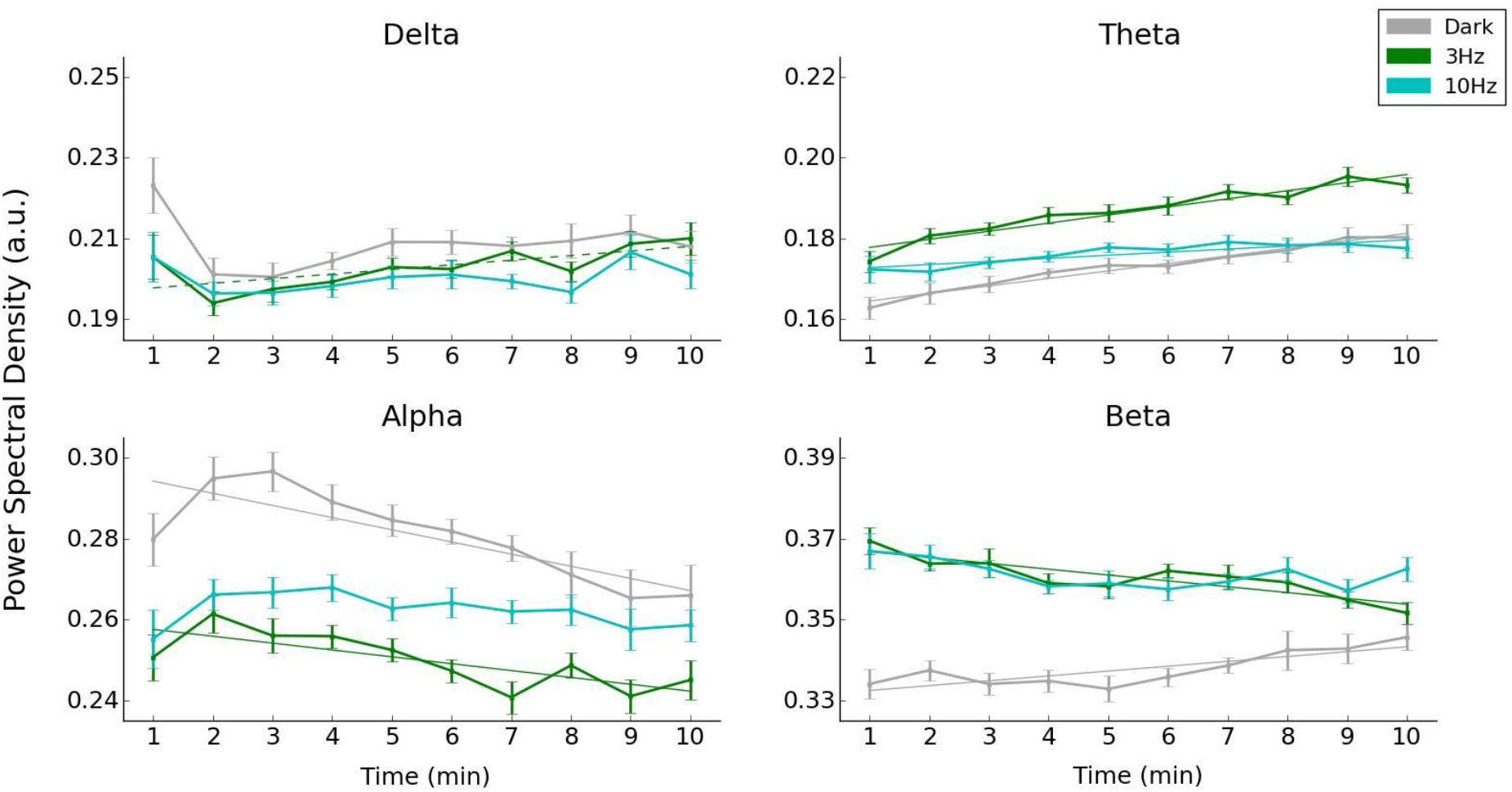
Mean power spectral density for each condition and frequency band: delta (1-4 Hz), theta (4-8 Hz), alpha (8-13 Hz) and beta (13-30 Hz). Each point represents a non-overlapping 1-minute segment of data, across a total of 10 minutes. Dashed lines indicate least-squares regression *p*-values between 0.01 and 0.05, solid lines indicate *p* < 0.01, uncorrected. Units are power spectral density, measured in arbitrary units (a.u.), normalised by the sum of the spectral power of all frequency bands, resulting in each frequency band representing a percentage of the total spectral power. Error bars are normalised within-subjects^42^.

Finally, in additional exploratory analyses we tested if LZs and each normalised spectral power band are reflecting distinct features of the data. We computed the Pearson correlation coefficients r between all measures, on the single participant averages (averaged across channels and time) and for each condition (Fig 8). We only display correlations where r > 0.5, in order to highlight strong correlations. We observed a strong correlation across conditions between LZs and beta (r = 0.804, *p_bon_ =* .0003, Dark, r = 0.809, *p_bon_ =* .0003, 3 Hz, r = 0.816, *p_bon_ =* .0002, 10 Hz). We also note that beta power and LZs displayed similarities in their temporal pattern of results (Figs 5 and 7) i.e. a decreasing trend for 3 Hz and 10 Hz and increasing trend for Dark. Investigating the correlation between these two measures over time, using a Fisher’s z-transformation of r, revealed a strong correlation between beta and LZs for all conditions (Dark (r = 0.70, *p_bon_* = .001), 3Hz (r = 0.73, *p_bon_* < .001) and 10 Hz (r = 0.74, *p_bon_* < .001)). Suggesting that changes in beta power may be a critical feature of the power spectrum driving changes in LZs between conditions. Examining other correlations in the data we found that spectral power between alpha and delta was negatively correlated (r = −0.841, *p_bon_* < .0001, Dark, r = −0.743, *p_bon_* = .003, 3 Hz, and r = −0.81, *p_bon_ =* .0003, 10Hz). For 3 Hz only, we found a negative correlation between LZs and alpha (r = −0.688, *p_bon_* = .01). While for 10 Hz only, we found a negative correlation between beta and theta (r = −0.679, *p_bon_* = .01).

**Figure 8.**
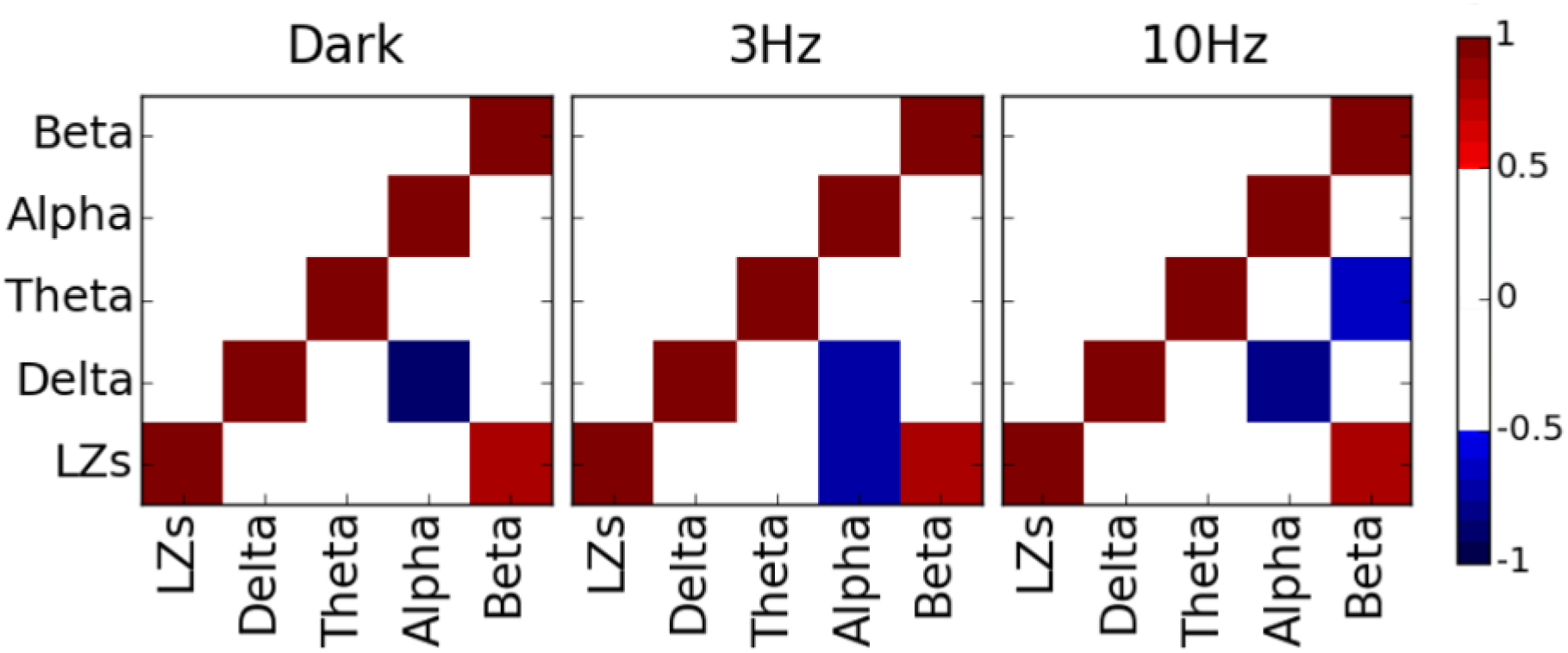
For each condition, a matrix shows the Pearson correlation coefficients, r, across participants, between each pair of measures. For clarity, the upper triangular portion of the symmetric correlation matrices and entries with |r| < 0.5 are omitted (set to white) to highlight strong correlations only. All correlations are Bonferroni corrected for multiple comparisons.

## Discussion

Using stroboscopic stimulation, we induced non-pharmacological ASC while measuring the diversity (Lempel-Ziv) and power of EEG signals. Stroboscopic stimulation at 3 Hz and 10 Hz, with eyes closed, gave rise to dissociable experiences that included marked increases in intensity and phenomenological ratings (ASCQ), which were accompanied by simple, and in some cases, complex visual hallucinations (CVH). These alterations in experience were accompanied by significant increases in spontaneous EEG signal diversity for both 3 Hz and 10 Hz stimulation (above a baseline of wakeful rest), a similar pattern of results to those found with psychedelic compounds (LSD, psilocybin, ketamine)^8^. By utilising two surrogate data sets, we found that changes in both the power spectrum and the phase spectrum contributed to the increases in EEG signal diversity. Examining changes in the power spectrum during stroboscopic stimulation, we found a significant reduction in alpha power during 3 Hz stimulation, (relative to Dark, Fig 5), in which participants also reported the majority of CVH occurred. We note that similar alterations in spectral power have been observed during psychedelic states^8, 13, 14^. Together, our results support the hypothesis that neural signal diversity (LZs) provides a neurophysiological signature of the diversity of subjective experiences that are associated with different global states of consciousness.

Characterising the phenomenological effects of stroboscopic stimulation at 3 Hz and 10 Hz via subjective reports (intensity and ASCQ ratings), we found that both frequencies led to marked increases in the intensity and range (diversity) of experiences compared to Dark (Fig S8). While there were some commonalities in experience between 3 Hz and 10 Hz, they also differed across multiple ASCQ dimensions, as well as in the overall subjective intensity of experience (section 4.1). 10 Hz was characterized by high intensity, simple SIVHs, with some commonalities in experience to a psilocybin induced ASC. In contrast, the 3 Hz condition was characterized by a relatively low intensity, simple SIVHs, fewer similarities to a psilocybin induced ASC, but interestingly, in addition, was associated with the occurrence of complex visual hallucinations (CVH). Together with the between-condition differences in the power spectrum, our results reveal that stroboscopic stimulation at differing frequencies is capable of producing substantially different states of consciousness.

Our main finding of reliable changes in LZs, during non-pharmacological altered states of consciousness, provides further support that, in addition to indexing alterations in consciousness level, such as sleep stages or anaesthetic level^6, 7^, LZs shows sensitivity to the diversity of experience. Specifically, we suggest that increases in LZs during ASC may reflect an increase in the temporal diversity of experiences or percepts. A key feature of each conscious experience is that subjectively it is experienced as being unique and distinct from other experiences. This observation has led to the intuitive assumption that each conscious experience must be reflected by a specific pattern of cortical activity, with two similar experiences evoking relatively similar patterns of cortical activity. For instance, as I glance across my desk, the pattern of cortical activity produced by looking at my coffee mug in one moment should be fairly similar to looking at the keyboard in the next moment. By contrast, during psychedelic ASC, perception appears to be less continuous in terms of the temporal rate of change of conscious experience^10, 13^, i.e. an experience in one moment may be very different to an experience in the next moment, such as during non-veridical hallucinations. Therefore, during psychedelic, and possibly stroboscopic ASC, an increase in the temporal diversity of experience may be reflected by a wider range of distinct patterns of cortical activity in a given time window compared to the normal waking state. Measures of EEG signal diversity that are sensitive to changes in global brain dynamics, would therefore naturally reflect the increased temporal diversity of experience associated with these states. This suggestion is in line with the previously reported enhanced repertoire of brain dynamical states found via fMRI during a psilocybin-induced psychedelic state^44^. There are also many converging lines of evidence supporting the notion that ASC, such as the psychedelic state, are associated with ‘increasing the bandwidth of perceptual experience’ (for more detail see ^3^). For example, a large cluster analysis of ASC questionnaire responses taken from 327 psilocybin sessions^11^, found that the dimension ‘everything around me was happening so fast that I no longer could follow what was going on’ was more likely to be scored highly following psilocybin, possibly reflecting an increase in the temporal diversity of experience. Our suggestion is also compatible with efforts to more accurately describe states of consciousness^3^. Compared to the normal wakeful state, both psychedelic and stroboscopic ASC represent substantially different states of consciousness, yet neither is well described as representing a higher or lower ‘level’ of consciousness. This observation has led some to suggest that states of consciousness are best described as regions in a multidimensional space^3, 9^. The findings of the present study, along with our previous application of LZs to psychedelic states^8^, demonstrate that increases in signal diversity occur alongside marked increases in the intensity and range/diversity of subjective experiences (Figs 4 and S8). If, as we suggest, EEG signal diversity reflects the temporal diversity of phenomenal experience, then it would provide a useful dimension for characterising global states of consciousness, indexing both brain dynamics and the diversity of experience, within such a multidimensional space.

In apparent contrast to our primary findings linking elevated LZs to SIVH, when investigating variations in LZs over each 10 min session, in an exploratory analysis, we found a clear decreasing linear trend during both 3 Hz and 10 Hz conditions, with a clear increasing trend during the Dark condition (Fig 5). As far as we are aware, previous research has not explored the variability of LZs over time during the waking state. If as we suggest and LZs is sensitive to the diversity of experiences, then the changes in LZs with time across conditions may reflect changes in the temporal diversity of experience. For example, during stroboscopic conditions the initial experience of geometric patterns and colourful kaleidoscopic imagery may have increased the temporal diversity of experience, which according to our theory would lead to an enhanced repertoire of brain states, and therefore more diverse neural activity and subsequent higher LZs scores. The decreasing trend in LZs for stroboscopic conditions may reflect a transition between the rapidly changing experiences to more stable complex visual phenomena. In contrast, during the Dark condition, with no visual input the diversity of participants mind wandering may have increased over the 10 min session causing the increasing trend in LZs scores. It is important to recall that the only continuous subjective measure we recorded was of intensity (analogue slider scores), not range or diversity. We found, for all conditions, an increase in intensity over time (Fig S8). Therefore, the changes in LZs over time cannot be explained by changes in subjective intensity, suggesting that LZs is not sensitive to the intensity of experience. Future experiments are needed that investigate the effect of selectively manipulating the diversity of subjective experience on LZs, and to track the evolution of subjective diversity over time, to shed further light on these issues.

In our previous application of LZs to source-localised MEG data obtained during psychedelic states^8^, we demonstrated, using phase-randomised surrogate data, that the observed increases in signal diversity could not be entirely explained by changes in spectral power. However, we did not investigate the influence of the phase spectra on signal diversity during psychedelic ASC. Here, we were able to separate the contributions of both the power and phase spectra to LZs using two types of surrogate data. Using EqualPhase surrogate data, we found a strong condition-specific influence of the power spectrum on LZs (Fig 4), but we also found that changes in power could not entirely account for the changes in LZs. With the EqualPower surrogate data, we also found a significant increase in signal diversity for 3 Hz and 10 Hz relative to baseline (Dark), showing a specific contribution of phase information to changes in LZs. Together, these surrogate data analyses reveal that differences in both the power and phase spectra contributed to the increase in signal diversity during stroboscopic stimulation relative to baseline (Dark). Given that an EEG signal can be completely described in terms of its spectral power and phase, what then does LZs add to the description of the signal? One response is that while changes in the amplitude of a particular frequency band are important when describing states of consciousness, for example increased delta power (1-4 Hz) during NREM, there is no single spectral band that displays the same sensitivity to states of consciousness in an analogous manner to LZs. LZs captures the spontaneous signal diversity of brain activity - with the theoretically-motivated link to phenomenological diversity - whereas there is no such obvious relationship with any single spectral band. Therefore, changes in the power and phase spectrum should be viewed as separate signatures of brain dynamics compared to measures of neural signal diversity, such as LZs.

Examining the changes in LZs scores between conditions we note the similarities between the results of the original and EqualPhase data (Fig. 4). These results suggest that changes in signal diversity associated with stroboscopic ASC were strongly influenced by the power spectrum, in a similar manner to psychedelic ASC^8^. What specific alterations in the power spectra could be contributing to these changes in signal diversity? The most pronounced spectral alteration, which appears to be consistent across different psychedelic compounds (LSD, psilocybin and ketamine) is the marked decrease in alpha power^8, 13, 14, 16^.

Alpha oscillations (8 –12 Hz) have been associated with the excitability of cortical sensory networks, with high levels of alpha being associated with increased cortical inhibition and therefore reduced excitability of the visual network^45-47^. During psychedelic ASC, a reduction in alpha power has been suggested to reflect disinhibition, increasing cortical excitability^13, 16^. A reduction in normal inhibitory processes may engender a shift away from externally stimulus-driven toward internally-driven information processing, and in doing so increase the likelihood of visual hallucinations^13^. In line with this view, several computational models have shown that an increase in the excitability of the visual network (in the absence of visual input) destabilizes spontaneous neuronal activity, creating elementary (simulated) visual hallucinations^19, 48, 49^. Together, these results provide a plausible model of the origins of psychedelic visual hallucinations. During psychedelic ASC a reduction in alpha power marks decreased cortical inhibition, facilitating the spread of spontaneous internally generated patterns of neural excitation over the visual cortex, leading to the experience of visual hallucinations^13, 16, 48^.

Following from this model, decreases in alpha power during psychedelic ASC have also been linked to the formation of complex visual hallucinations. For example, the magnitude of the reduction in alpha power during psychedelic ASC (LSD) has been shown to predict the extent of both simple and complex visual hallucinations, as well as more profound changes in consciousness, such as ego-dissolution^16^. These results are also supported by findings from other psychedelic compounds, such as psilocybin, which show that the decrease in alpha power correlates with the formation of visual hallucinations^13^.

Interestingly, in the present study, we also observed a pronounced decrease in alpha power during 3 Hz stroboscopic stimulation (relative to Dark, Fig 6), in which participants reported the majority of complex visual hallucinations to occur. Similar to psychedelic ASC, we speculate that a reduction in alpha power due to stroboscopic stimulation reflect disinhibition within the visual network, although the mechanisms underlying how stroboscopic stimulation would lead to a increased disinhibition and reduced alpha power admittedly remain unclear. However, computational models investigating the effects of stroboscopic stimulation on visual cortex postulate that simple visual hallucinations (SIVH, geometric patterns etc.) may be caused by dynamic instabilities within primary visual cortex^48-50^. In particular, Rule et al., ^50^ found that stroboscopic stimulation affected lateral inhibition within a model of visual cortex, producing spatial instabilities within the network that underlie the experience of (simulated) geometric hallucinatory patterns. Interestingly, clinical conditions in which both simple and complex visual hallucinations are common have also implicated alterations in lateral inhibition^25, 51^.

To summarise, our results, in combination with modelling and psychedelic investigations of visual hallucinations during ASC, support a decrease in alpha power as being associated with the formation of complex visual hallucinations during ASC, likely through increased disinhibition within visual cortical networks.

Continuing our exploratory analysis, investigating other features of the power spectra that could have contributed to changes in signal diversity, we found a strong positive correlation between LZs and the beta frequency band (13-30 Hz) across all conditions (Fig 8). Our previous application of LZs to psychedelic MEG data did not find a similar relationship between these two measures (see Fig 2 in ^8^). Examining the temporal evolution of LZs and beta power, we found a similar pattern of condition-specific changes over the course of the 10 min session (Fig 4 and 6) i.e. a decreasing trend for 3 Hz and 10 Hz, and an increasing trend for Dark. Among a variety of roles, beta-band activity has been associated with the maintenance of current cognitive state, ensuring the dominance of top-down signals that override the effect of unexpected external events^52, 53^. In the current study, we speculate that the decreasing trend in beta power during stroboscopic conditions may have facilitated the influence of bottom-up stroboscopic signals, allowing spatial instabilities to spread within primary visual cortex, leading to the experience of geometric hallucinatory patterns. Further experiments, perhaps involving exogenous manipulations of beta-band activity, are needed to substantiate these ideas.

Examining the verbal descriptions of visual experience and the overall phenomenological profile (ASCQ) produced by stroboscopic stimulation we note certain similarities to a psilocybin-induced psychedelic state. Firstly, consistent with previous reports^18, 21-23, 25, 31^, during 3 Hz and 10 Hz stimulation all participants reported ‘simple’ geometric patterns and colourful kaleidoscopic imagery. These types of simple visual hallucinations, such as ‘three-dimensional patterns’ or ‘colourful flickering kaleidoscopic imagery’ are also reported during the psychedelic state^12^,^36, 54^. Secondly, psychedelic ASC are often reported to include CVH such as life-like objects or scenes^12^,^36, 37, 54^. Interestingly, we found that following 3 Hz stimulation the majority of participants also reported CVH, comprising of detailed perceptions of faces and scenes, which were described as being highly vivid, disjointed and bizarre visual experiences, which were subjectively close to veridical perception (section 4.2). Additionally, while not explored explicitly in the present study, stroboscopic stimulation has also been shown to induce a range of dissociative phenomena relating to the sense of identity/self ^31, 32^, again similar to those reported during the psychedelic state^33, 34, 36, 37^.

The parallels between experiences reported during stroboscopic and psychedelic ASC motivate future research to explore stroboscopic stimulation as a non-pharmacological method of inducing an ASC that is qualitatively similar in some respects to the psychedelic state. However, we recognise that stroboscopic or psychedelic ASC are not fully characterised by the questionnaire and report measures used. Psychedelic and stroboscopic ASC differ in many respects, for example in terms of the profundity of experience. Psychedelic ASC are reported as being similar to life-changing spiritual or mystical experiences^9, 55^, which can lead to enduring increases in prosocial attitudes and behaviours as well as healthy psychological functioning^56^. Additionally, psychedelic experiences typically occur with eyes open, with the inherent interplay between the hallucinatory content and normal perception. Therefore, while we report that stroboscopic stimulation shows some similarities to psychedelic ASC, we recognise that these states of consciousness also differ in many respects.

While CVH are a common feature of psychedelic ASC^12, 36, 37, 54^, they have not been previously reported to occur as the result of stroboscopic stimulation^18, 21-23, 57^. What factors in our experiment may account for their emergence? There are a number of possibilities, first, the stroboscope used in this study (Lucia N^0^03) is considerably brighter (single LED power = 700 lm, x8 LEDs = 5600 lm) than those used in previous studies that have examined SIVH (for example^18^, single LED power = 80 lm power, x3 = 240 lm, as a reference, a 100 W light bulb produces approximately 1600 lm). Stimulus luminance is known to increase the magnitude of neural evoked responses at early stages of visual processing^58^. The high luminance used in this study may therefore have led to stronger effects on inhibitory processing on visual cortex, which may account for the emergence of CVH.

It is also possible that the CVH we observed were the result of the specific stimulation frequencies, with the majority of CVH occurring during 3 Hz stimulation. Until now, the subjective effects of 3 Hz stroboscopic stimulation has rarely been examined due to it inducing (overall) less intense visual experiences compared to frequencies closer to dominant alpha (8-12 Hz)^21, 25^. Interestingly, in our study experiences of stroboscopically-induced CVH during the 3 Hz condition were unrelated to self-reported intensity of experience (participants rated 3 Hz to be less intense than 10 Hz (Fig 3)). Previous work has shown that the content of simple, but not complex, visual hallucinations (i.e. radial, grid or spiral patterns), can be modulated by the specific stroboscopic stimulation frequency^18, 24^. As previously mentioned, stroboscopic stimulation at 3 Hz caused entrainment of this frequency, SSVEP, along with its harmonics, leading to a significant increase in theta power for 3 Hz relative to the other conditions (Fig. 6). This may have led to an increase in the relative proportion of lower frequencies such as delta and theta compared to normal. Increases in theta power during ASC have often been associated with the hypnagogic state^59, 60^. It is interesting to note that in many respects, participants’ phenomenological reports of CVH during 3 Hz stimulation resembled the ‘fantastic visual phenomena’ sometimes reported during hypnagogic states^61^. A defining feature of a hypnagogic state is the report of discrete, highly vivid, visual experiences, which lack a narrative or overarching theme^59, 61, 62^. Similarly, participants in this study reported that CVH lacked a dream-like narrative and were described as comprising of detailed perceptions of objects or scenes, yet were rated as being surprisingly close to veridical perception. Similar to hypnagogic experiences, all participants were aware of the unreal nature of their CVH^59, 61, 62^. Previous research investigating hypnagogic states and their EEG correlates during drowsiness and sleep onset have found that the spontaneous, transient, visual imagery connected with hypnagogic states were associated with decreased alpha and/or increased theta power^59, 60, 63^. We note that only during the 3 Hz condition did we observe reports of CVH that resembled those of hypnagogic experiences, in combination with a ‘hypnagogic-like’ spectral pattern of results i.e. a significant decrease in alpha and increase in theta power, relative to both Dark (Fig 6).

In summary, it is likely that specific parameters of our setup, LED luminance and specific stimulation frequency contributed to the occurrence of CVH in this study. Stroboscopic CVH display some commonalities to both experiential reports and alterations in power spectrum observed during hypnagogic hallucinations. Since we did not anticipate the appearance of CVH in this study, further research is needed to examine the precise conditions under which CVH arise through stroboscopic stimulation.

The resurgence of scientific interest into psychedelic drugs has been partially driven by renewed interest in their clinical therapeutic potential. However, psychedelic drugs are not without risks, with as many as 1 in 25 regular users reporting ongoing long-term visual and dissociative disturbances^64^. The ability of stroboscopic stimulation to induce altered phenomenology with marked similarities to psychedelic ASC, along with similar changes in neural activity (increased signal diversity, reduced alpha power), suggests it may provide an alternative to or adjunct method alongside psychedelic therapy.

## Conclusion

By combining stroboscopic stimulation with EEG, we found that stimulation at 3 Hz and 10 Hz generates subjectively striking changes in experience (as measured by the ASCQ), which were accompanied by increases in EEG signal diversity (as measured by Lempel-Ziv complexity (LZs)) compared to wakeful rest. These subjective reports and increases in signal diversity show similarities to those observed during psychedelic states engendered by LSD, ketamine, or psilocybin^41^, indicating that spontaneous signal diversity provides a robust signature of ASC. Using surrogate data, we demonstrated that increases in signal diversity under stroboscopic stimulation depended on changes in both the power spectrum and the phase spectrum of the underlying EEG. Although stroboscopic and psychedelic ASC differ in many respects, our findings of substantial changes in experience, along with both ‘simple’ and ‘complex’ visual phenomena, demonstrate that stroboscopic stimulation offers a powerful non-pharmacological means of inducing ASC, as well as providing a possible adjunct to psychedelic therapies. Overall, our results provide further evidence that EEG signal diversity reflects the diversity of subjective experiences that are associated with different states of consciousness.

## Acknowledgments

We thank Dirk Proeckl, Engelbert Winkler and Light Attendance gmbh for helpful discussions and loaning us Lucia N^0^03. We also thank Satohiro Tajima for useful discussions on data analysis, and Maxine Sherman for helpful comments on a draft version of this manuscript. DJS, AC and AKS are grateful to the Dr. Mortimer and Theresa Sackler Foundation which supports the Sackler Centre for Consciousness Science. Michael Schartner is supported by the University of Geneva, Department for Fundamental Neurosciences. AKS is also grateful to the Canadian Institute for Advanced Research (CIFAR) Azrieli Programme on Mind, Brain, and Consciousness.

## Author contributions

DJS, MS and AKS designed the study and wrote the paper. DJS, BA, FS and AYC collected the data. DJS, MS, BA and FS analysed the data. All authors reviewed and approved the final manuscript.

