## Supplementary Information for "Increased spontaneous EEG signal diversity during stroboscopically-induced altered states of consciousness"

David J. Schwartzman [a,b*], Michael Schartner [b,c], Benjamin B. Ador [d], Francesca Simonelli [e], Acer Y.-C. Chang [b,f], Anil K. Seth [a,b]

[a] Department of Informatics, University of Sussex, Brighton BN1 9QJ, United Kingdom

[b] Sackler Centre for Consciousness Science, University of Sussex, Brighton BN1 9QJ, United Kingdom

[c] Département des Neurosciences Fondamentales, Université de Genève, 1206 Genève

[d] Master de Biologie, École Normale Supérieure de Lyon, Université Claude Bernard Lyon I, Université de Lyon, 69342 Lyon Cedex 07, France

[e] IMT School for Advanced Studies Lucca, Lucca, Italy

[f] Department of Neuroinformatics, Araya Brain Imaging, Tokyo, Japan

*

#### Behavioural Pilot

In order to identify two phenomenologically distinct stroboscopic stimulation frequencies, which caused substantial changes in experiential content compared to wakeful rest, we conducted a behavioural pilot study.

##### Participants

Eleven healthy students (mean age of 21.4, seven female) of the University of Sussex completed the behavioural pilot. Participants provided informed consent before taking part and received £10 or course credits as compensation for their time. The experiment was approved by the University of Sussex ethics committee.

##### Methods

The behavioural pilot used the same equipment and design as the main experiment unless otherwise stated. Previous investigations into the effects of stroboscopic stimulation have found that the most vivid 'simple' visual experiences are reported using a stimulation frequency close to the dominant alpha frequency (8-12 Hz), with stimulation frequencies below 5 Hz or above 30 Hz causing subjectively weaker visual experiences [1-5]. Based on these findings we chose to investigate four stroboscopic frequencies, three outside the alpha frequency range (3 Hz, 5 Hz and 40 Hz), in an attempt to find a frequency that produced mild experiences, and one (10 Hz) close to the dominant alpha frequency. Participants were exposed to 5-minute sessions at each stimulation frequency, with the order of the stimulation frequencies randomized for each participant. Following each session participants answered the Altered States of Consciousness Questionnaire (ASCQ adapted from [6, 7], see Fig3A for dimensions and questions).

#### Results

Comparing responses for all ASCQ questions, we found that the 3 Hz caused the weakest shifts in experience, scoring the lowest on all ASCQ dimensions except for the 'Space', 'Merge', and 'Control' dimensions (Fig S1). Based on [8], we then considered the mean score, referred to as 'TOTAL', over all ASCQ questions to index the overall intensity of each stroboscopic state, comparing the results of each condition using paired *t*-tests (Bonferroni corrected). The results revealed that the 3 Hz condition displayed the weakest experiences, showing significantly lower 'TOTAL' responses compared to 5Hz ($t(10) = -5.5$, $p < .001$), 10 Hz ($t(10) = -9.2$, $p < .001$) and the 40 Hz conditions ($t(10) = -2.6$, $p < .001$). In contrast, stroboscopic stimulation at 10 Hz produced the largest 'TOTAL' ASCQ score across participants, as well as higher overall scores for the following dimensions: 'Arousal', 'Intensity', 'Ego', 'Imagery', 'Float', 'Merge', 'Mood', 'Muddle', 'Patterns', 'Space', 'Spirit', 'Strange' and 'Vivid' (Figure S1).

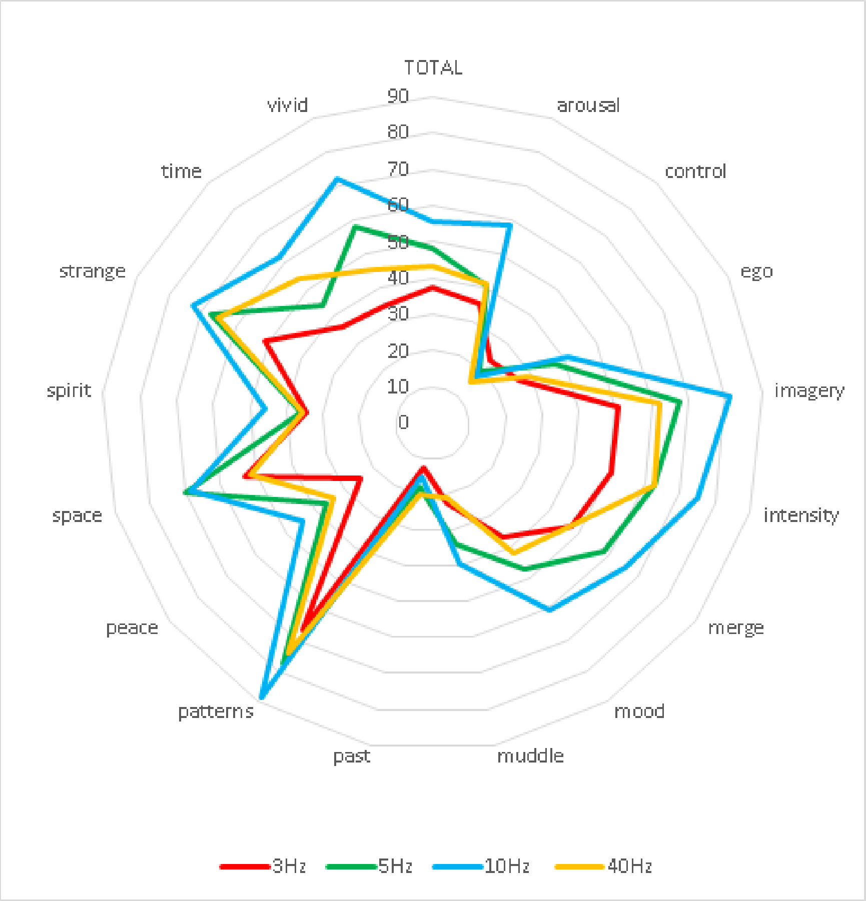


**Figure S1**. Radar plot of absolute ASCQ responses, averaged across participants. Responses for each stroboscopic stimulation frequency are shown: 3 Hz ('red'), 5 Hz ('green'), 10 Hz ('blue'), 40 Hz ('orange'). For ASCQ questions see Fig 3A.

To examine if 3 Hz and 10 Hz stimulation produced dissociable states, we compared average ASCQ responses, finding that the overall intensity 'TOTAL', ($p < .003$) as well as 'Vivid' ($p = .026$), 'Arousal' , ($p = .005$), 'Merge' ($p = .04$), 'Mood' ($p = 0.005$), 'Imagery' ($p = .006$), 'Patterns' ($p = .007$), 'Strange' ($p = .004$) and 'Time' ($p = .04$) dimensions were significantly lower for 3 Hz compared to 10 Hz. Based on these results we chose 3 Hz and 10 Hz as the stimulation frequencies for use in the main experiment as they produced the weakest and most intense experiences, respectively, while also differing significantly across multiple ASCQ dimensions.

### Supplementary Analyses

#### Topographic distribution of condition-specific changes in LZs

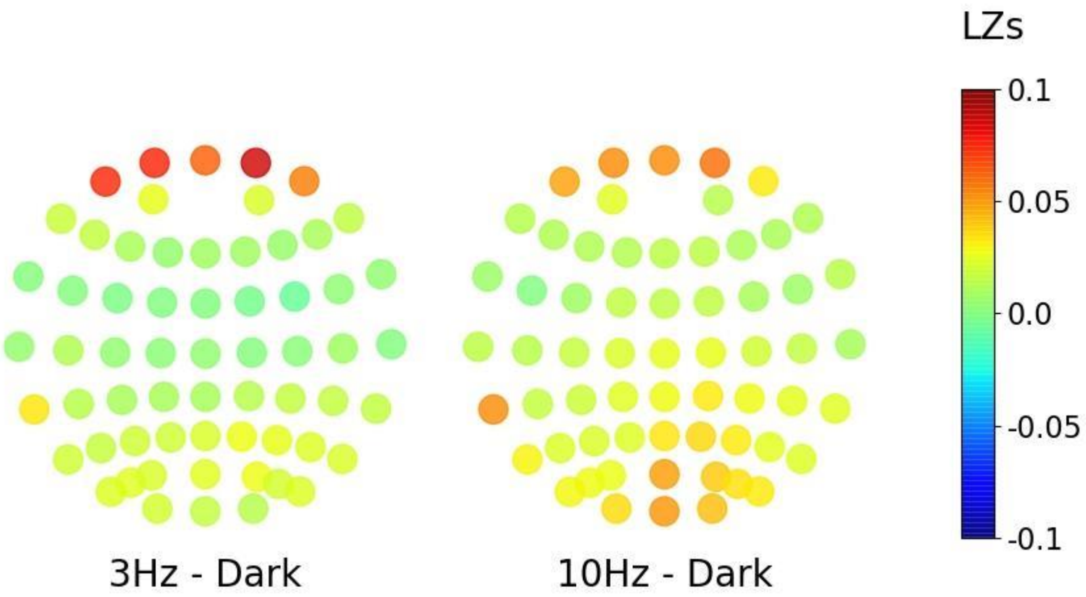


**Figure S2.** Topographic plots showing the difference in mean LZs scores for original data, top of each plot shows frontal electrodes. For each electrode we show the difference of LZs scores for each stroboscopic condition minus the baseline condition (Dark) i.e. 3 Hz-Dark and 10 Hz-Dark.

#### Topographic distribution of condition-specific changes in the power spectrum

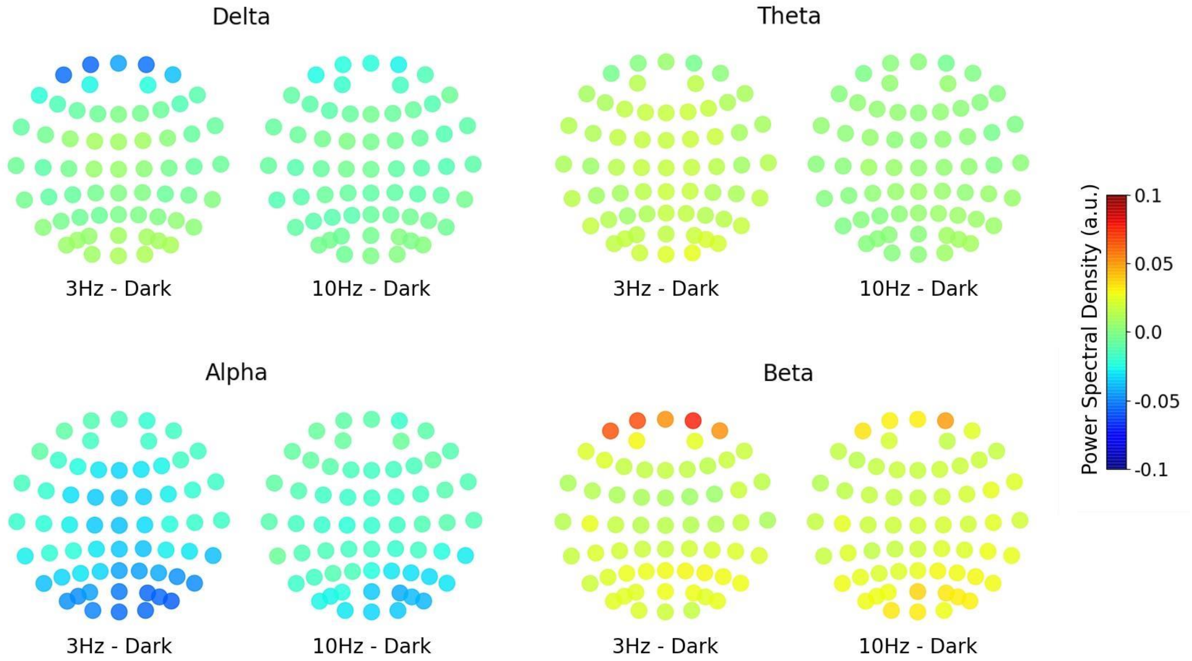


**Figure S3.** Topographic plots displaying the difference in average power spectral density for each frequency band: delta (1-4 Hz), theta (4-8 Hz), alpha (8-13 Hz) and beta (13-30 Hz) for each stroboscopic condition minus Dark i.e. 3 Hz-Dark and 10 Hz-Dark. Units are power spectral density, measured in arbitrary units (a.u.), normalised by the sum of the spectral power of all frequency bands, resulting in each frequency band representing a percentage of the total spectral power (i.e., relative not absolute power).

#### Steady-state analyses – notch filtered EEG data

To exclude the possibility that steady state visual evoked potentials (SSVEPs) contributed to the condition specific alterations in LZs, we applied periodic Butterworth notch filters (width = 0.5Hz, order = 4 or 5 (3 Hz and 10 Hz, respectively) at the appropriate stroboscopic stimulation frequencies (3 Hz or 10 Hz) prior to further analysis. This process led to the creation of two additional data sets: 'Notch 3 Hz' and 'Notch 10 Hz'. To investigate the effects of the notch-filters on the SSVEPs we applied a one-dimensional discrete Fourier Transform to the full 10 minute data for each condition, participant and electrode and then averaged across these dimensions (Fig S4). As can be seen in Fig S4 the notch filters successfully removed the majority of the SSVEPs for both stimulation frequencies.

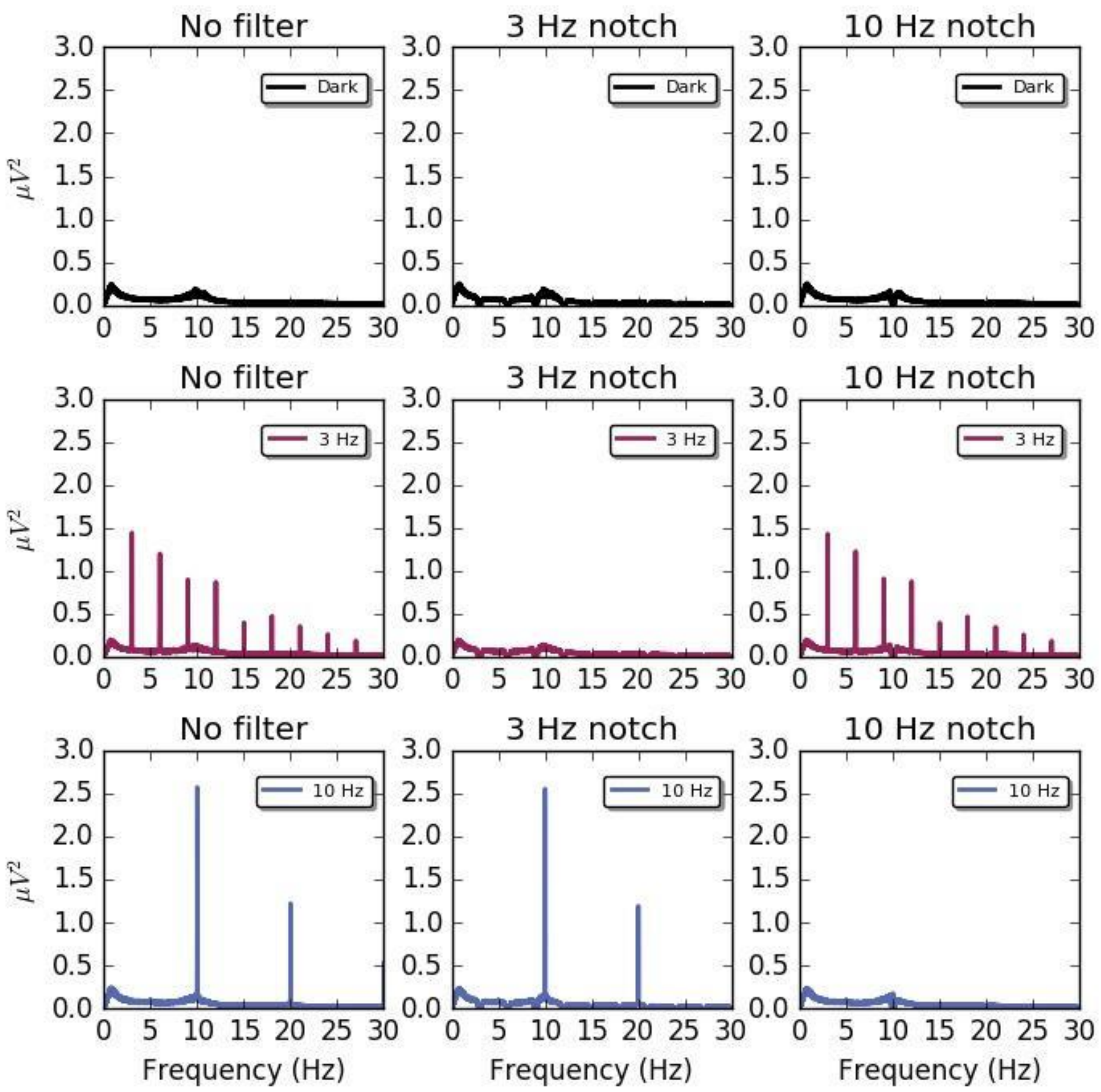


**Figure S4**. Average power spectrum of each 10-minute session for the original data (no filter) or with a 3 Hz or 10 Hz Butterworth notch filter applied. Data was averaged across all participants and electrodes. The top panels display the spectral power of the Dark condition and with a 3 Hz (from 3 to 30 Hz in steps of 3 Hz) and 10 Hz (from 10 to 30 Hz in steps of 10 Hz) notch filters applied. The middle left panel shows the effects of entrainment of the 3 Hz stroboscopic signal on the power spectrum. As can be seen in central panel the effects of entrainment (both at driving frequency and harmonics) are successfully removed by the notch filter. Similarly, the bottom left panel displays the power spectrum corresponding to 10 Hz stroboscopic stimulation and the rightmost panel following a 10Hz notch filter.

#### Effects of SSVEP on signal diversity

Having confirmed that the notch filters successfully removed the SSVEP, we next computed LZs for the notch-filtered data. Note that in order to allow for a fair comparison between original and notch-filtered data, we also applied each notch-filter to the Dark condition. As in the main experiment, we found increases in signal diversity for both 3 Hz (Notch 3 Hz: $t(18) = -6.02$, $p < .0001$; Notch 10 Hz: $t(18) = -3.72$, $p < .0001$) and 10 Hz notch-filtered data (Notch 3 Hz: $t(18) = -6.79$, $p < .0001$; Notch 10 Hz: $t(18) = -4.45$, $p < .001$) relative to Dark. However, the difference in LZs between 3 Hz and 10 Hz conditions was no longer evident for the notch-filtered data (Fig S5).

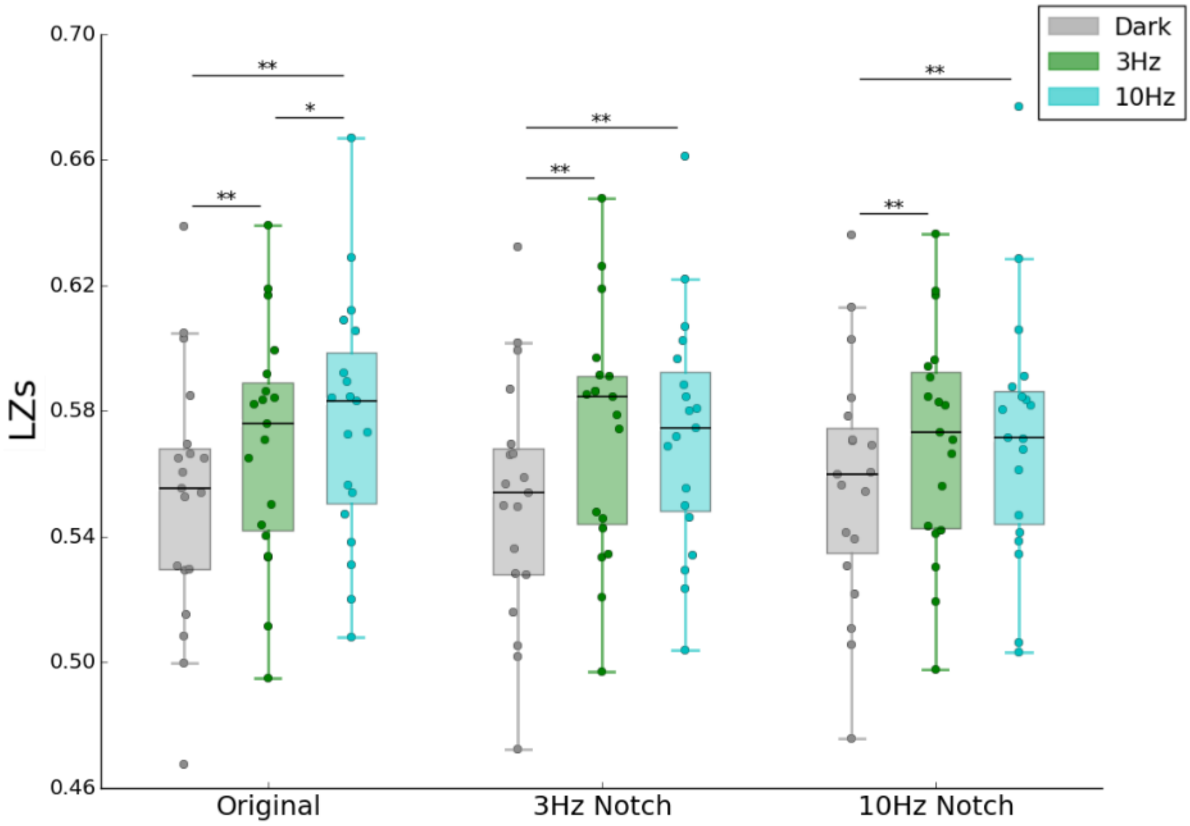


**Figure S5**. Mean signal diversity scores (LZs) for each condition for Original, 3 Hz and 10 Hz Butterworth notch filtered data. Original data shown for comparison. 3Hz Notch: mean signal diversity scores (LZs) across conditions following a 3 Hz notch filter (3 to 30 Hz in steps of 3 Hz). 10Hz Notch: mean signal diversity scores (LZs) across conditions following a 10 Hz notch filter (10 to 30 Hz in steps of 10 Hz). The box extends from the lower to upper quartile values of the data, with a horizontal line at the median. Whiskers display the range of the data (excluding outliers). Circles show individual participant results. ** indicates a *p*-value < 0.01, * indicates a *p*-value >0.01 < 0.05, obtained using paired sample *t*-tests (Bonferroni corrected).

#### Effect of SSVEP on power spectra

To investigate the effects of SSVEPs on the endogenous spectral profile, we computed the same spectral comparisons as in the main experiment, this time using the 3 Hz and 10 Hz notch-filtered data. As can be seen in Fig S6, the results confirmed that SSVEPs due to stroboscopic stimulation at 3 Hz and 10 Hz, indeed influenced the spectral profile of the data. However, as can be seen in Fig S6, examining the spectral profile of each frequency band for the original data, in which the SSVEP are still present, all of the strongly significant results ($p = < 0.01$) associated with each frequency band and condition are still evident in the notch-filtered data. Suggesting that while there was an effect of SSVEPs on the power spectrum, overall this contribution was minimal compared to endogenous condition-specific alterations in the power spectrum.

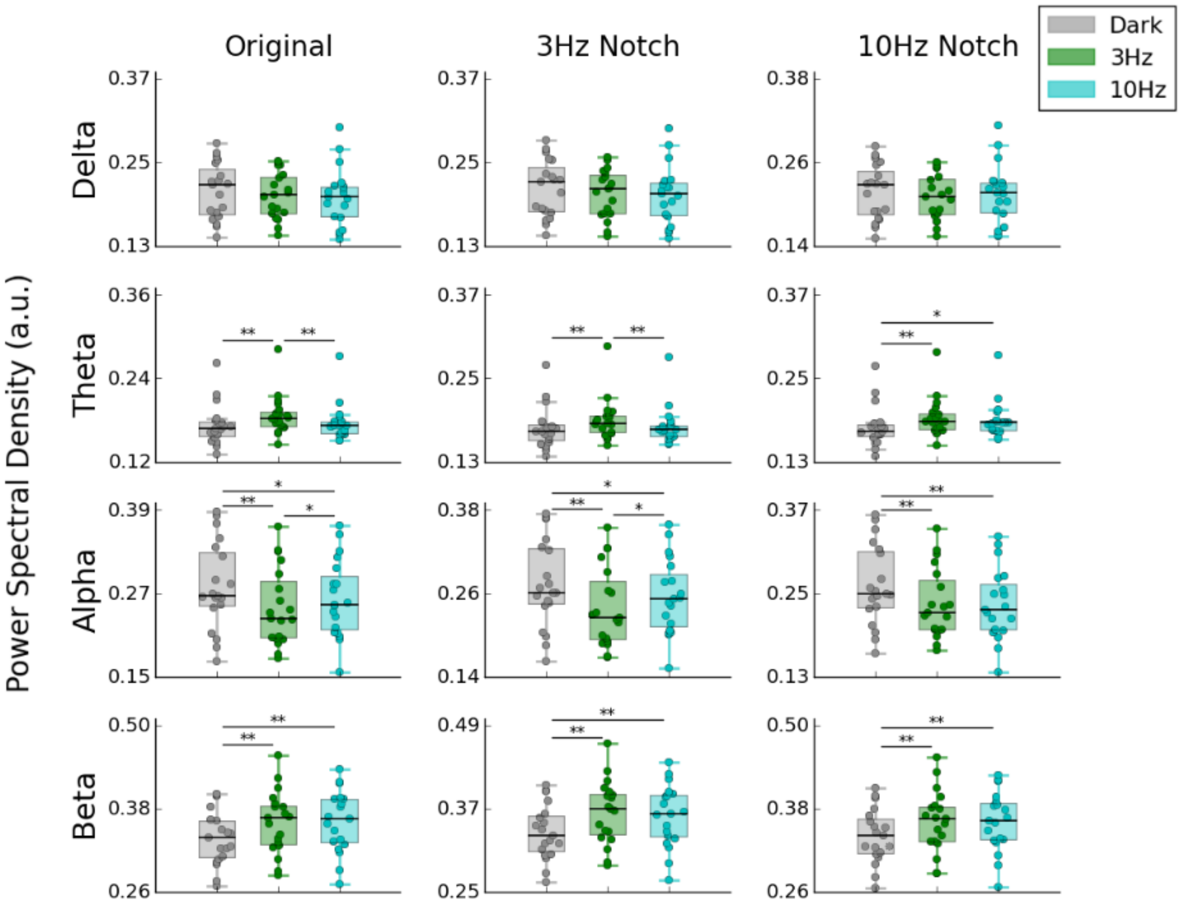


**Figure S6**. Average power spectral density of original and notch-filtered EEG data for each condition and frequency band: delta (1-4 Hz), theta (4-8 Hz), alpha (8-13 Hz) and beta (13-30 Hz). Original data shown for comparison. Each box extends from the lower to upper quartile values of the data, with a horizontal line at the median. Whiskers display the range of the data (excluding outliers). ** indicates a $p$-value < 0.01, * indicates a $p$-value >0.01 and <0.05, obtained using paired sample $t$-

tests (Bonferroni corrected). Units are power spectral density, measured in arbitrary units (a.u.), normalised by the sum of the spectral power of all frequency bands, resulting in each frequency band representing a percentage of the total spectral power.

#### Single participant signal diversity scores

To examine the robustness of our main signal diversity result, we examined condition-specific changes in LZs at the single participant level. As can be seen in Fig S7, the broad pattern of LZs results observed at the group level i.e. increased signal diversity scores for stroboscopic compared to Dark conditions, also holds for the majority of participants.

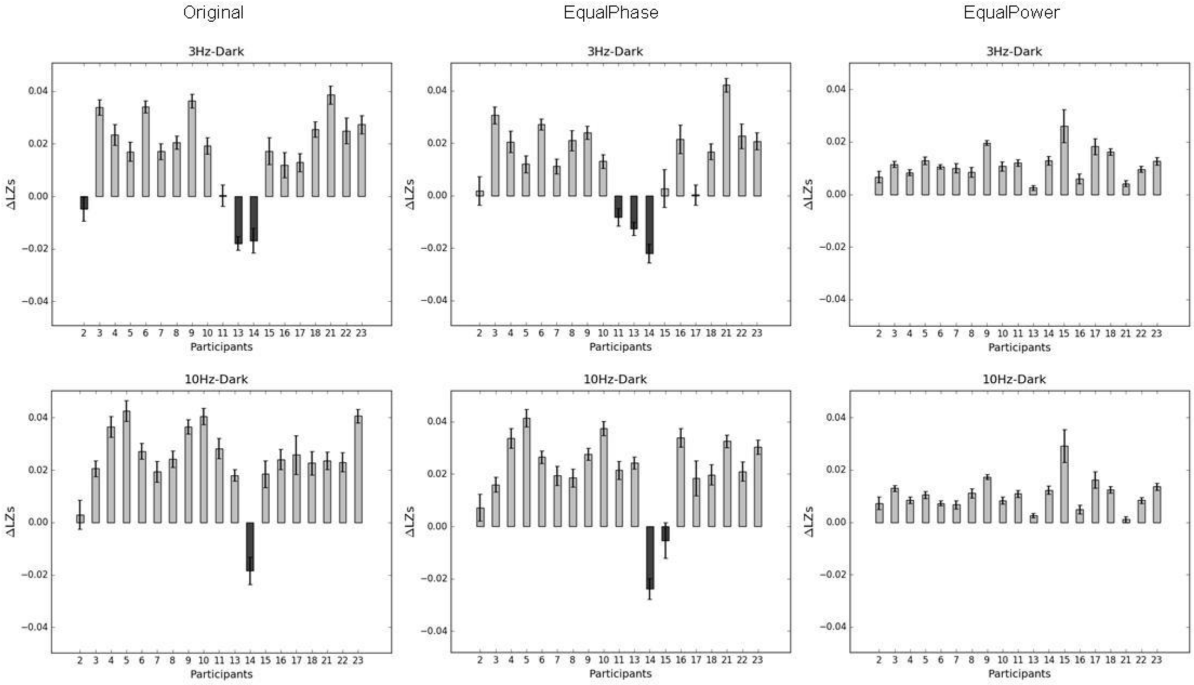


**Figure S7**. Difference in mean signal diversity (LZs) scores across conditions, 3 Hz-Dark (top panels) and 10 Hz-Dark (bottom panels) for each participant, using original and surrogate data. Within-subjects error bars were calculated using the method described in O'Brien and Cousineau (2015).

### Mean signal diversity and intensity rating scores across time

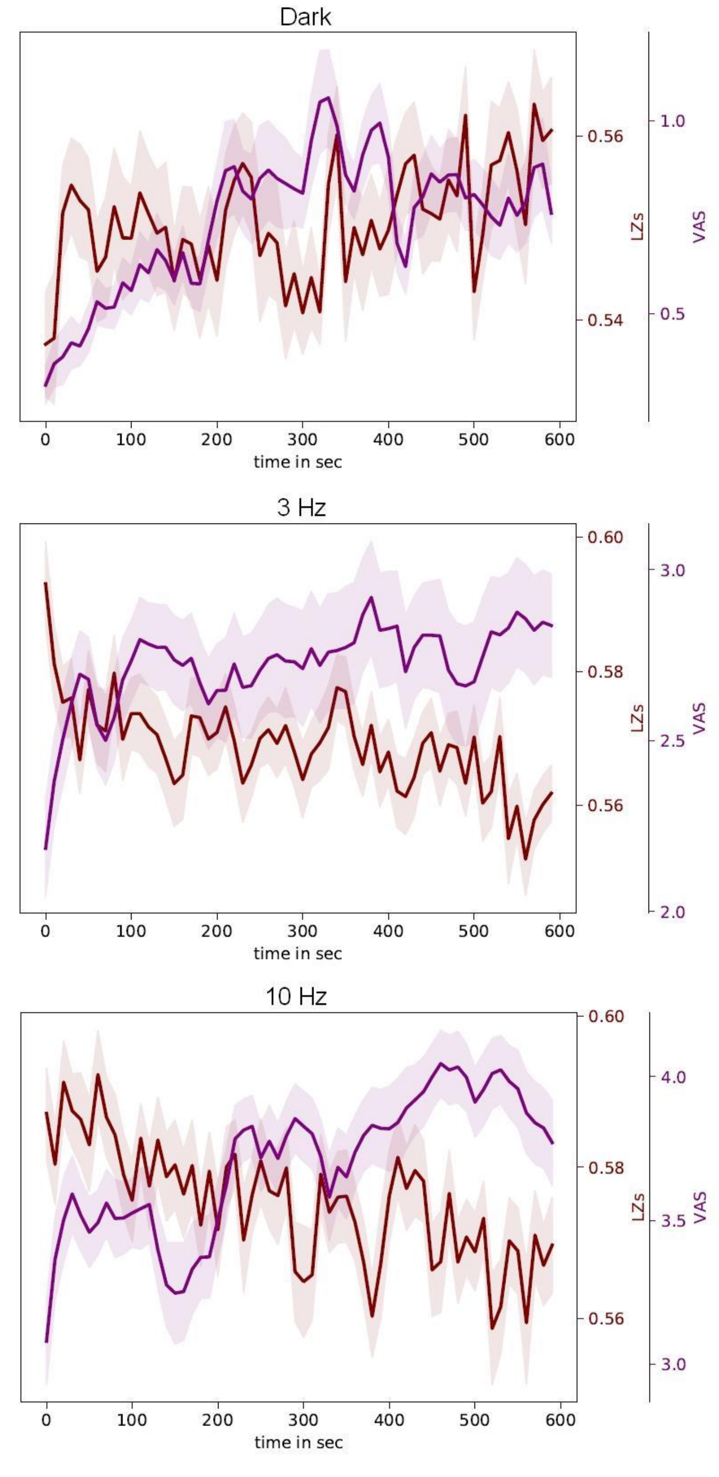


**Figure S8**. Mean signal diversity (LZs) across time for the original data and intensity ratings given throughout each session via the visual analogue slider (VAS) for each condition. LZs was computed for each participant, channel and non-overlapping 10 sec segment. Red lines display the average LZs value across channels and participants. Purple lines display VAS responses averaged across participants and each non-overlapping 10 sec segment of data. The error band displays the standard error across participants. As can be seen, the intensity of experience increased across the 10 minute session for all conditions, with the largest increase in intensity being observed for 10 Hz.
